# Parasitic flatworms infecting thorny skate, *Amblyraja radiata:* infection by the monogeneans *Acanthocotyle verrilli* and *Rajonchocotyle emarginata* in Svalbard

**DOI:** 10.1101/2020.11.19.389767

**Authors:** Raquel Hermans, Maarten P. M. Vanhove, Oleg Ditrich, Tomáš Tyml, Milan Gelnar, Tom Artois, Nikol Kmentová

## Abstract

Parasite diversity above the Arctic circle remains understudied even for commercially valuable host taxa. Thorny skate, *Amblyraja radiata,* is a common bycatch species with a growing commercial value. Its natural range covers both sides of the North Atlantic including the Arctic zone. Svalbard is a Norwegian archipelago located on the northwest corner of the Barents Shelf which sustains a spectacular species diversity. So far, several monogenean species have been reported infecting thorny skate across the Atlantic Ocean. In the present study, we intend to fill in the knowledge gap on monogenean parasites infecting thorny skate in the northern part of its range and thus indirectly assess the connectivity between the thorny skate populations off the Svalbard coast and from previously studied locations. 46 monogenean individuals were recovered from 11 specimens of thorny skate. Following morphological and molecular assessment, two species of monogeneans, *Acanthocotyle verrilli* and *Rajonchocotyle emarginata,* were identified. The results serve as the northernmost record for both parasite genera and the first record of monogenean species off Svalbard. Detailed morphometric evaluation revealed a relatively high level of morphological variation in *A. verrilli* compared to its congeners. Phylogenetic reconstruction placed *A. verrilli* in a well-supported clade with *A. imo.* Our study also suggests high diagnostic significance of sclerotised structures in the identification of *Rajonchocotyle.* Even though the occurrence of two directly transmitted parasite species supports the previously suggested long-distance migration of *A. radiata*, future studies employing highly variable genetic markers are needed to assess the ongoing and historical migration patterns.

**Highlights:**

- First record of monogenean species in Svalbard
- Northernmost record for representatives of Acanthocotylidae and Hexabothriidae
- Transatlantic occurrence of parasites supports connectivity of thorny skate populations

## 1. Introduction

Thorny skate (Chondrichthyes, Rajidae) is a common bycatch species with growing commercial value. It prefers shallow coastal waters with muddy or sandy substrate [1] and temperatures from −1.4°C to 16°C [2]. Increased fishing effort during the last decades severely impacted the overall biomass of this skate species characterized by low fecundity, slow growth rate, and late maturity [3,4]. Its known geographic distribution ranges from South Carolina in the Western part of the Atlantic Ocean to Greenland and the North Sea in the East and Svalbard in the Arctic zone [5–11].

Svalbard is an archipelago in the Arctic Ocean located on the northwest corner of the Barents Shelf. The Arctic Ocean is the smallest of all oceans with a mean depth of 1361 m and a total area of approximately 10 million km^2^ [12,13]. It consists of four abyssal plains surrounded by continental shelves comprising c. 50% of the total area [13]. The northern and western margins of the Barents Shelf end in the continental slope down to the Polar Ocean Basin and the oceanic Norwegian Greenland Sea, respectively [14]. The wide range of habitats on continental shelves sustains a spectacular biodiversity in this marine ecosystem [15] and harbours species of Atlantic and Pacific affinities due to ancient connections. However, continental shelves were an important migration barrier especially to shallow-water organisms [16] between the Arctic and adjacent oceans [17] during the last Pleistocene glacial/interglacial cycles. Overall, the inventory of biodiversity on Svalbard is far from complete because of the focus of most studies on its west coast in view of the better accessibility of this region. Similar to terrestrial habitats, the inventory of marine biodiversity off Svalbard’s coast has been limited and biased by sampling techniques (e.g., pelagic trawls) or towards certain taxonomic groups such as Crustacea and Mollusca [18,19].

In general, parasite biodiversity in the Arctic is mostly understudied and many species remain unknown including fish parasites [20–23]. Data about parasite fauna of *Amblyraja radiata* from the North Atlantic and Arctic regions of its distribution are missing. Scientific exploration of the marine parasite fauna in this part of the world has been mainly concentrated on Franz Josef Land and heteroxenous parasite taxa [22]. Recently, Murzina et al. [24]reported on the parasite fauna of *Leptoclinus maculatus* (Perciformes, Stichaeidae) (Fries 1838), being the first record of parasitic flatworms (Trematoda) at the Svalbard coast. Globally, there are almost 1500 parasite species described from 900 elasmobranch species to date [25]. Helminth infections of the thorny skate were reported worldwide (see Table 1). So far, representatives of two monogenean families and three species have been reported. Monogenea is a group of parasitic flatworms (Neodermata, Platyhelminthes) with worldwide occurrence and a mostly ectoparasitic life-style. They are primarily parasites of fish characterised by a direct life-cycle and predominantly narrow host-specificity [26,27]. The basic division is between members of blood feeding Polyopisthocotylea and epithelial feeding Monopisthocotylea, representatives of both of which have been reported to infect the thorny skate [28,29]. The northernmost distribution of a parasitic flatworm infecting *A. radiata* was *Acanthocotyle verilli* Gotto, 1899 recorded off Tromsø, Norway [30],see Table 1). Hence, flatworm infections on *A. radiata* have never been recorded above the Arctic Circle.

**Table 1:**
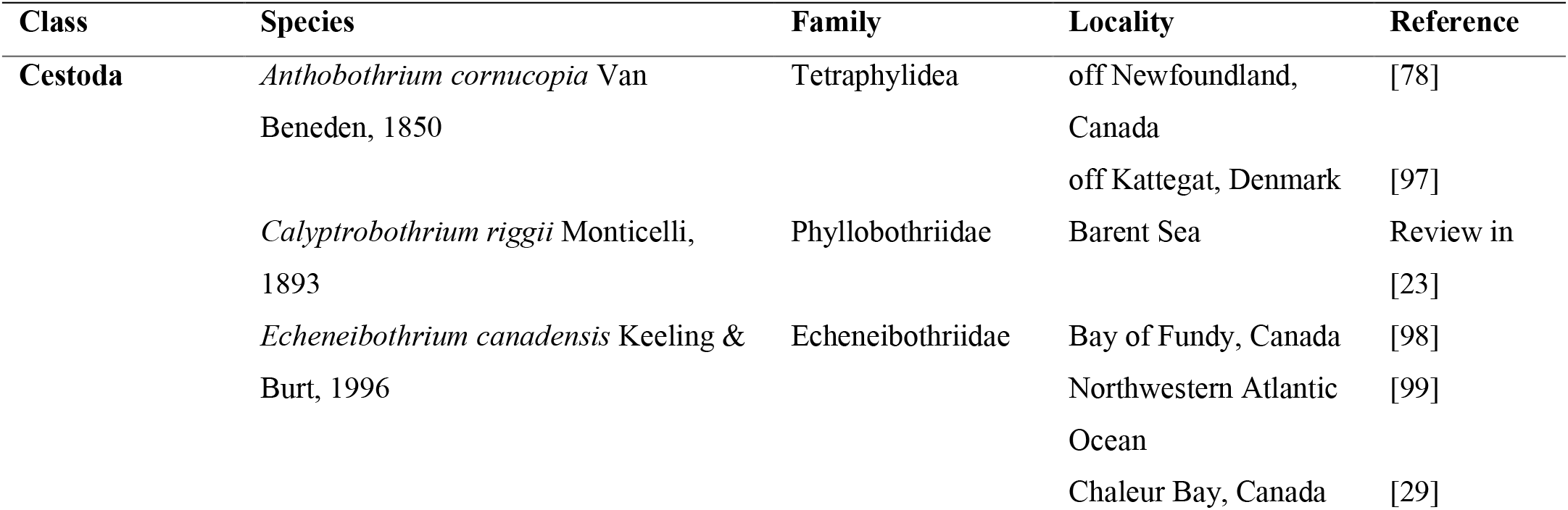
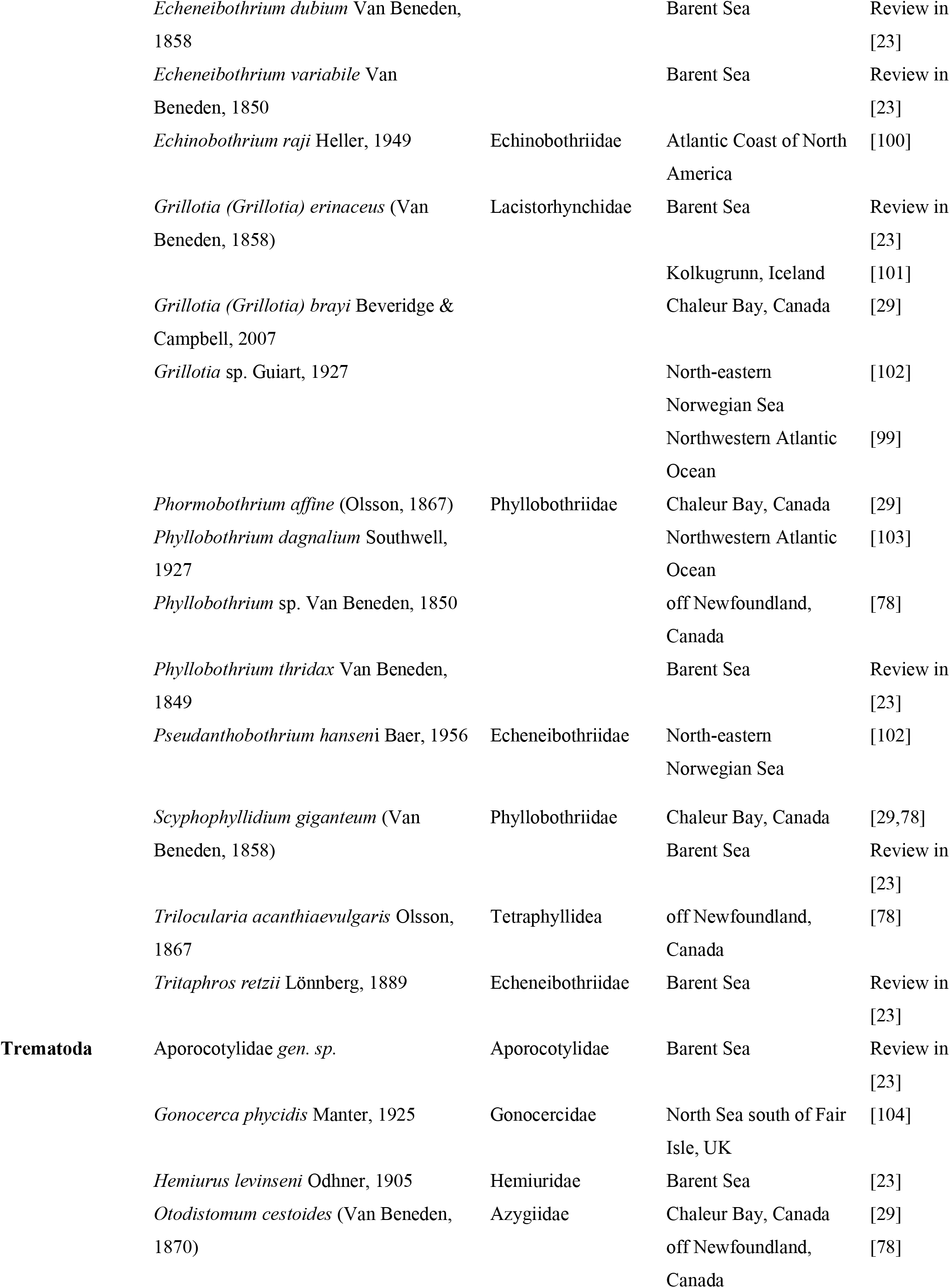
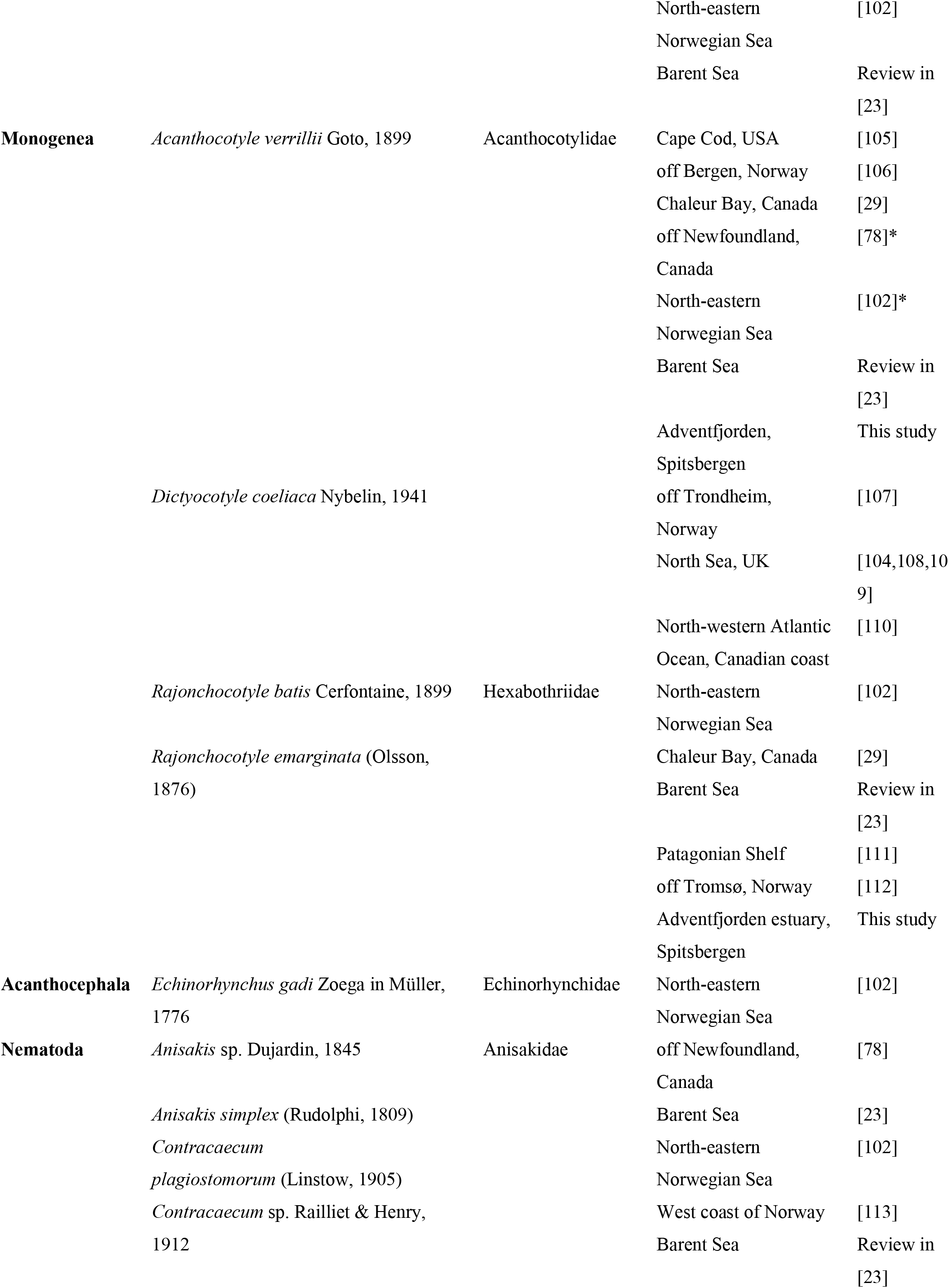
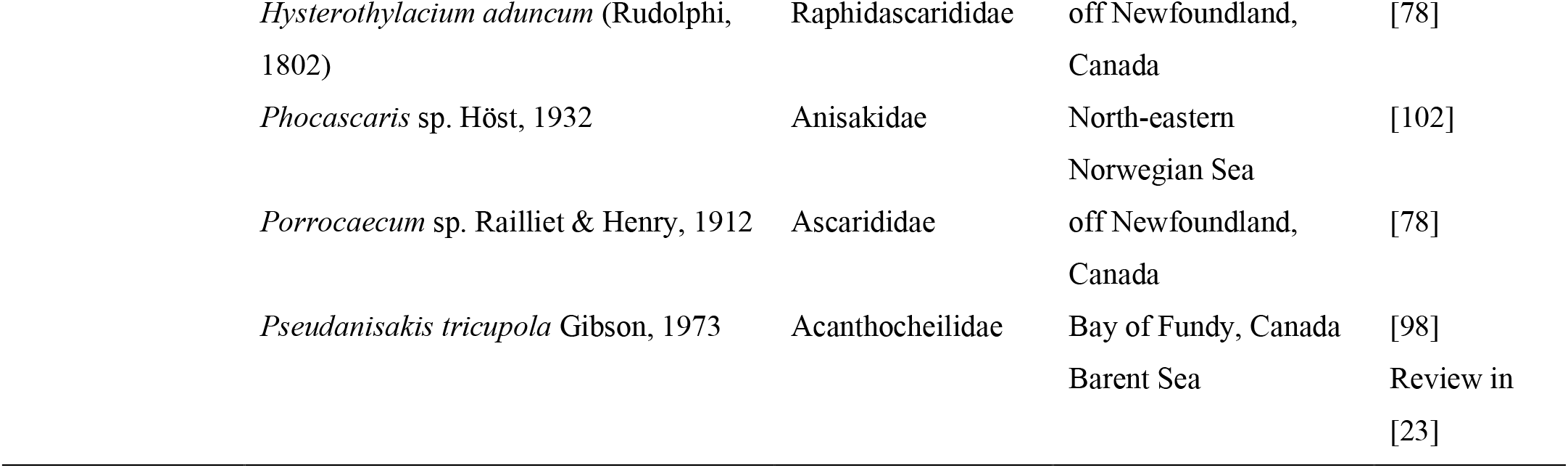
List of helminth species reported from *Amblyraja radiata* with taxonomic designation and locality of the report.

In the present study, we intend to fill in this gap of knowledge in the distribution of monogenean parasites infecting thorny skate, *A. radiata*, at the northern part of its known range, in Spitsbergen, Svalbard.

## 2. Material and Methods

### 2.1 Host collection

In total, eleven specimens of *A. radiata* were examined for the presence of monogenean individuals during a field expedition in Spitsbergen, Svalbard, July 2016 organised by the Centre for Polar Ecology (University of South Bohemia, Czech Republic). Host specimens were caught in the Adventfjorden near Hotellneset, Spitsbergen, Svalbard (78°15’18”N, 15°30’58”E) using benthic gill nets at a depth of 30–40 m and immediately transported to the laboratory in seawater containers. Prior to dissection and subsequent examination, skates were euthanised by overdosing with tricaine methane sulphonate (Sigma-Aldrich, Darmstadt, Germany).

### 2.2. Parasite collection and morphological examination

The fins, gills and nasal cavity were examined for the presence of monogeneans. Monogenean individuals were transferred with a needle and mounted on slides using a solution of glycerine ammonium picrate (GAP). Selected specimens were kept in 99% ethanol and subsequently stained using acetocarmine combined with Gomori trichrome, cleared with clove oil and mounted in Canada balsam. Two species were found in this study, *Acanthocotyle verilli* and *Rajonchocotyle emarginata.* Infection parameters per parasite species namely prevalence (percentage of infected hosts), infection intensity (mean number of monogenean individuals per infected host) and abundance (mean number of monogenean individuals per examined host) were calculated following Ergens and Lom [31]. In total, 13 and 21 morphological characters including hard and soft parts following Kearn et al. [32] and Bullard and Dippenaar (2003), respectively, were measured and photographed using a Leica DM 2500 LED microscope (Leica Microsystems, Wetzlar, Germany) and the software LasX v3.6.0. Voucher specimens are deposited in the collection of the Research Group Zoology: Biodiversity and Toxicology at Hasselt University in Diepenbeek, Belgium (HU) under the following accession numbers: xx-xx. Type specimens from the Helminthological Collection of the South Australian Museum, South Australia, Australia (AHC) and the National Museum of Natural History of the Smithsonian Institution, Washington, USA (USNM), were examined for comparative purposes: *Acanthocotyle atacamensis* Ñacari, Sepulveda, Escribano & Oliva, 2019 – 1 paratype (1 slide) USNM 1480281*; A. gurgesiella* Ñacari, Sepulveda, Escribano & Oliva, 2018 – 1 paratype (1 slide) USNM 1422089*; A. imo* Ñacari, Sepulveda, Escribano & Oliva, 2019 – 1 paratype (1 slide) USNM 1480278*; A. lobianchi* Monticelli, 1888 – 2 vouchers (2 slides) AHC 36231, 36232; *A. pacifica* Bonham & Guberlet, 1938 – 2 paratypes (2 slides) USNM 1321942; *A. pugetensis* Bonham & Guberlet, 1938 – 2 paratypes (2 slides) USNM 1321940; *A. verrilli* – 1 paratype (1 slide) USNM 135051; *A. urolophi* Kearn, Whittington, Chisholm & Evans-Gowing, 2016 – 4 paratypes (4 slides) AHC 36222, 36223, 36224, 36225; *Branchotenthes octohamatus* Glennon, Chisholm & Whittington, 2005 – 2 paratypes (2 slides) AHC28769, 28770; *B. robinoverstreeti* Bullard & Dippenaar, 2003 – 1 holotype (1 slide) USNM 1387687; *Callorhynchocotyle callorhynchi* (Manter, 1955) – 2 paratype (2 slides) AHC29747; *C. amatoi* Boeger, Kritsky & Pereira, 1989 – 1 paratype (2 slides) AHC29749; *Erpocotyle antarctica* (Hughes, 1928) – 2 paratypes (2 slides) AHC29725; *E. somniosi* – 4 paratype (4 slides) USNM 1349221; *Heteronchocotyle gymnurae* Neifar, Euzet & Ben Hassine, 2001 – 1 paratype (1 slide) USNM 1385030; *Paraheteronchocotyle amazonensis* Mayes, Brooks & Thorson, 1981 – 2 paratypes (2 slides) USNM 1372658; *Rajonchocotyle emarginata* Olsson, 1876 – 1 paratype (1 slide) USNM 1337399; *R. laevis* Price, 1942 – 1 syntype (1 slide) USNM1337422; *R. wehri* Price, 1942 – 4 paratypes (4 slides) USNM 1337421; *Squalonchocotyle borealis* (Van Beneden, 1853) Cerfontaine, 1899 – 4 paratypes (4 slides) USNM 1349221; *S. callorhynchi* – 1 paratype, 1 holotype (2 slides) USNM 1338129; *S. impristi* – 1 holotype (1 slide) USNM 1338749. Selected specimens of both collected species were drawn using a drawing tube and afterwards edited using the software GIMP v2.10.20. Interspecific morphological differences and the level of intraspecific phenotypic variability were evaluated using measurements relative to the total length of the parasite’s body because of the correlation between morphological variables and the total length [34]. Morphometric parameters where subsequently analysed by principal component analysis (PCA) in the R software package stats [35] and visualised using ggplot2 [36]. Only specimens mounted on slides with GAP were part of the analyses to avoid effects of the staining method on the results [37]. The following variables were used in PCA: (1) the ratios body width/total length (TL), (2) body length/TL, (3) pharynx length/TL, (4) pharynx width/TL, (5) diameter of the pseudohaptor/TL, (6) number of sclerite rows/TL, (7) testes maximum width/TL, (8) germarium length/TL and (9) germarium width/TL. Raw measurements are provided as Supplementary material (see Table S1&S2).

### 2.3 Scanning electron microscopy (SEM)

Following host examination, live monogenean individuals were fixed with hot 4% neutral buffered formaldehyde solution and transported to the Parasitology Laboratory of the Centre for Polar Ecology in České Budějovice, Czech Republic. Subsequently, samples were washed in 0.1 M phosphate buffer solution (three times for 15 minutes), post-fixed in 2% osmium tetroxide solution (for 60 minutes), washed again and dehydrated with an ascending acetone series (30%, 50%, 70%, 80%, 90%, 95%, 100%), each step for 15 minutes. Following dehydration, monogenean specimens were dried in liquid CO2 using a critical point method, placed on metal targets using double-stick tape, gold coated in a BAL-TEC SCD 050 sputter coater (Bal-Tec, Los Angeles, USA) and observed using a SEM JEOL JSM-7401F scanning electron microscope (JEOL, Tokio, Japan) at the Laboratory of Electron Microscopy, Institute of Parasitology, Biology Centre CAS in České Budějovice.

### 2.4 Molecular data generation

The posterior part of the body or complete specimens were used for genomic DNA isolation. The total genomic DNA was extracted using the Qiagen Blood and Tissue Isolation Kit (Qiagen, Hilden, Germany) following the manufacturer’s instructions. A portion of the large ribosomal subunit (28S rRNA) gene was amplified using the primer combination C1 (5’-ACCCGCTGAATTTAAGCAT-3’) and D2 (5’-TGGTCCGTGTTTCAAGAC-3’) [38]. Each PCR reaction contained 1.5 unit of *Taq* Polymerase (ThermoFisher Scientific, Waltham, USA), 1X buffer containing 0.1 mg/ml bovine serum albumin, 1.5 mM MgCl2, 200 mM dNTPs, 0.5 mM of each primer and 50 ng of genomic DNA in a total reaction volume of 30 μl under the following conditions: initial denaturation at 95 °C for 2 min followed by 39 cycles of 94 °C for 20 s, annealing at 58 °C for 30 s, extension at 72 °C for 1 min and 30 s, and a final extension step at 72 °C for 10 min. PCR products were purified using ExoSAP-IT (ThermoFisher Scientific, Waltham, USA) under the following conditions: 15 min at 37 °C and 15 min at 80 °C. Targeted DNA fragments were sequenced using the same primers as in the amplification reactions together with a Big Dye Chemistry Cycle Sequencing Kit v3.1 (ThermoFisher Scientific, Waltham, USA). Following clean up using the BigDye XTerminator Purification Kit (ThermoFisher Scientific, Waltham, USA), fragments were visualised on an ABI 3130 capillary sequencer (ThermoFisher Scientific, Waltham, USA). Electropherograms were visually inspected and assembled in MEGA7 [39]. The obtained sequences were deposited in NCBI GenBank under the accession numbers MW260310-12.

### 2.5 Phylogeny

Phylogenetic placement of collected monogenean species was inferred based on 28S rDNA at family level. Sequences generated during this study were aligned using MUSCLE [40] under default distance measures as implemented in MEGA v7 [39], together with 28S rRNA gene sequences of other *Acanthocotyle* spp. Selected representatives of related monogenean families (Bothitrematidae, Anoplodiscidae and Udonellidae) were used as outgroup [41]. Genetic distances among the species of *Acanthocotyle* were calculated as the pairwise difference (uncorrected p-distance) in MEGA v7 [39]. Poorly aligned and overly divergent regions were trimmed using Gblocks v0.91b [42] under the less strict flanking position option and allowing gap positions within the blocks. The final alignment consisted of 825 bp. The most appropriate evolutionary model, the HKY + Γ model [43], was selected based on the Bayesian information criterion in jModelTest v2 [44]. Phylogenetic relationships were inferred under Bayesian inference (BI in MrBayes v3.2.0 [45]) based on two independent runs (20^5^ generations, sampled every 1,000th generation and with a burn-in of 10%). Parameter convergence and run stationarity were assessed in Tracer v1.6 [46]. Moreover, a maximum likelihood (ML) search was performed in RAxML v8.2.12 with tree search conducted using RAxML’s standard tree search algorithm and bootstrap support calculated using the option with an automated number of replicates to obtain stable support values under the frequency stopping criterion [47]. Phylogenetic trees were edited in FigTree v1.4.2 (http://tree.bio.ed.ac.uk/software/figtree).

## 3. Results

In total, 37 and 9 monogenean individuals were found on fins and gills, respectively. Based on a detailed morphological examination, two species of monogeneans were identified.

To comply with the regulations set out in article 8.5 of the amended 2012 version of the International Code of Zoological Nomenclature [48], details of *Acanthocotyle verrilli* and *Rajonchocotyle ermaginata* have been submitted to ZooBank based on their respective original descriptions. For each taxon, the Life Science Identifier (LSID) is reported in the taxonomic summary.

### 3.1. *Acanthocotyle verrilli* Goto, 1899

Monogenea van Beneden, 1858

**Family** Acanthocotylidae Monticelli, 1903

**Genus** *Acanthocotyle* Monticelli, 1888

**Type-host**: *Amblyraja radiata* (Donovan, 1808) (Chondrichtyes, Rajidae)

**Other hosts:** *Bathyraja spinicauda* (Jensen, 1914) (Chondrichtyes, Arhynchobatidae); *Leucoraja erinacea* (Mitchill, 1825) (Chondrichtyes, Rajidae)

**Site on host:** Fins.

**Prevalence:** 72,7% (8 out of 11 infected)

**Intensity of infection:** 4,6 (1-12)

**Abundance:** 3,4 (0-12).

**Type-locality:** Cape Cod, USA

**Other localities:** Adventfjorden, Spitsbergen; Chaleur Bay, Canada; Coast of Maine, USA; off Bergen, Norway; off Newfoundland, Canada; North-eastern Norwegian Sea

**ZooBank registration:** The LSID for *Acanthocotyle verrilli* is urn:lsid:zoobank.org:act:FF7F506B-FB65-446C-9F91-255C45BF2E86.

#### 3.1.1. Remarks

In total, 13 morphological characters including soft body parts and sclerotised structures were measured (see Table 2, Figs. 1&2). Body elongate, circular pseudohaptor with radial rows of sclerites (26 – 34) covered by tegument (Fig. 2G). Pseudohaptor with 28 – 34 rows of sclerites, each row consists of 4 – 10 sclerites. Marginal valve of pseudohaptor with distinct fringe. The central part of the pseudohaptor, between the radial rows of sclerites, shows a depression indicating this part to be responsible for the attachment to the body (Fig 2E). True haptor with 16 marginal hooks located subterminally at the posterior margin of pseudohaptor (Fig. 2E). Hooks in the true haptor organised into one central pair and 14 hooks in a peripherical row, the latter pointing centrally (Fig. 2G) with shafts free of tegument (Fig. 2H). Pharynx globular. Three anterior adhesive lobes on each side of head with sense organs located at the side of each internal lobe (Fig. 2B). Accessory glands at the level of pharynx. Excretory bladders on each side, anterior to vitellarium field. Eyes absent. Testes (26 – 47) mainly rounded usually arranged in several (2 – 3) rows. Seminal vesicle unlobed, anterior part communicates with male genital opening via curved ejaculatory duct immediately posterior to pharynx. Male accessory gland reservoirs 2, adjacent to ejaculatory duct. Common opening at the level of intestinal bifurcation. Penis sclerite absent. Germarium immediately anterior to testes. Small uterine receptaculum seminis adjacent to germarium. Vagina absent. Germinal appendix not observed. Uterine pore opening on right side of the body, at the level of posterior part of pharynx. Eggs attached externally by an abopercular appendage (Fig. 2C). The egg operculum shows a pointed end (Fig. 2D). Vitelline follicles not discrete with lobed and overlapping margins, extending from level of germarium to near the posterior end of body proper.

**Table 2:** Meristic and morphometric data for *Acanthocotyle verrilli* ex *Amblyraja radiata* from Adventfjorden, Spitsbergen. Measurements are given in micrometers.

|  |  |
| --- | --- |
| Total body length | 1069–5855 (3541)<sup>a</sup> |
| Body proper length | 715–4688 (2757)<sup>b</sup> |
| Body width at the level of ovary | 330–925 (632)<sup>b</sup> |
| Pharynx |  |
| Length | 124–340 (233)<sup>b</sup> |
| Width | 77–413 (252)<sup>b</sup> |
| Pseudohaptor |  |
| Number radial rows of sclerites | 28–34 (mode=32)<sup>b</sup> |
| Length | 354–1167 (767)<sup>b</sup> |
| Width | 312–1112 (765)<sup>b</sup> |
| Testes |  |
| Testicular field length | 393–2583 (1522)<sup>c</sup> |
| Number | 18–47 (mode=34)<sup>d</sup> |
| Maximum width | 80–267 (173)<sup>c</sup> |
| Germarium |  |
| Length | 60–419 (215)<sup>c</sup> |
| Width | 98–347 (203)<sup>c</sup> |

Notes: <sup>a</sup>(n=28); <sup>b</sup>(n=29); <sup>c</sup>(n=26); <sup>d</sup>(n=22)

**Fig. 1:**
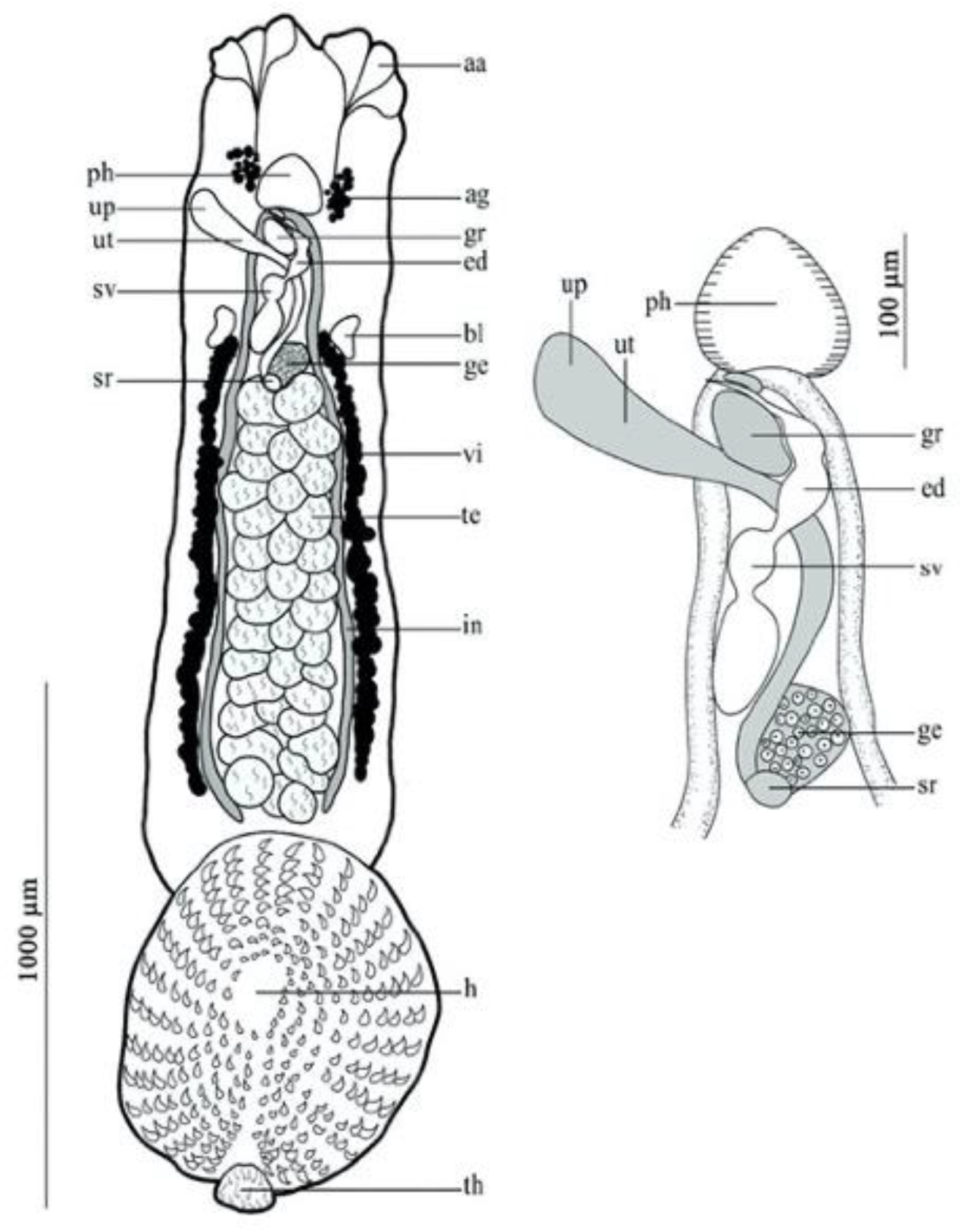
*Acanthocotyle verrilli* ex *Amblyraja radiata.* A) Ventral view of the whole body. B) Reproductive system. Abbreviations: aa, anterior adhesive lobes; ag, accessory glands; bl, excretory bladders; ed, ejaculatory duct; ge, germarium; gr, male accessory gland reservoir; h, haptor; in, intestine; ph, pharynx; sr, seminal receptacle; sv, bipartite seminal vesicle; te, testes; th, true haptor; up, uterine atrium; ut, uterus; vd, vas deferens; vt, vitelline duct; vi, vitelline.

**Fig. 2:**
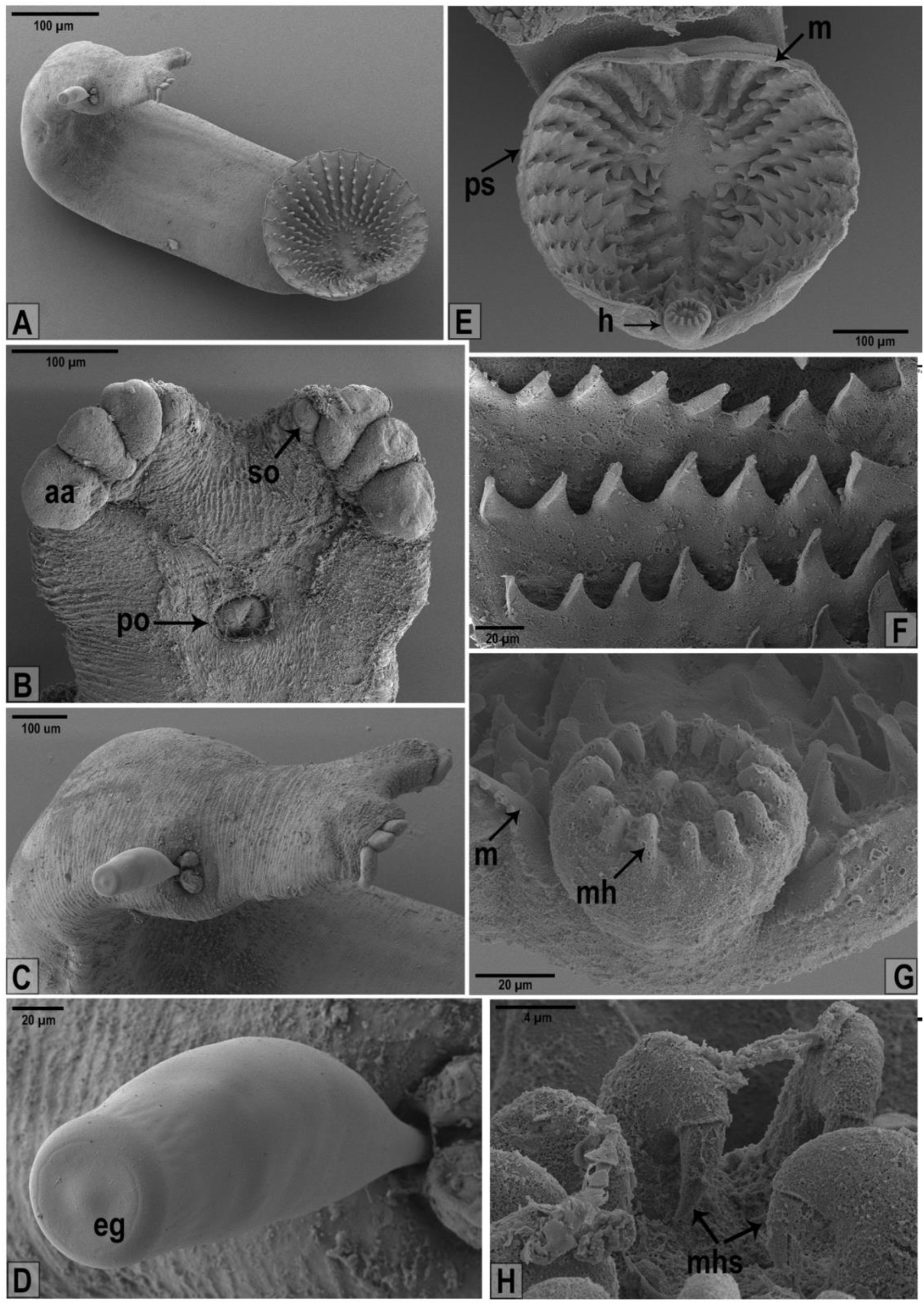
SEM observations of *Acanthocotyle verrilli.* A) Ventral view of the whole body, B) View of the anterior part of the body with adhesive lobes on each side, visible sense organ and pharynx opening. C) Uterus opening with an externally attached egg. D) Detailed apical view of an egg with abopercular appendage. E) Posterior part of the body formed by haptor and convex pseudohaptor, pseudohaptor with marginal valve. F) Detailed view of pseudophaptoral sclerites organised in rows. G) Detailed view of haptor located at the posterior margin of pseudohaptor armed with 14 peripheral and 2 central marginal hooks. H) aa – adhesive lobes, eg – egg operculum, h – haptor, m – marginal valve, mh – marginal hook, po – pharynx opening, mhs – marginal hooks sclerite, ps – pseudohaptor, so – sense organ.

#### 3.1.2. Differential diagnosis

Based on the recent revision of Acanthocotylidae published by Kearn et al. [32] combined with new species descriptions of Ñacari et al. [49,50], there are 12 species of *Acanthocotyle* currently considered valid: *Acanthocotyle atacamensis; Acanthocotyle elegans* Monticelli, 1890; *Acanthocotyle greeni* Macdonald & Llewellyn, 1980; *Acanthocotyle gurgesiella; Acanthocotyle imo*; *Acanthocotyle lobianchi*; *Acanthocotyle pacifica*; *Acanthocotyle patagonica* Kuznetsova, 1971; *Acanthocotyle pugetensis*; *Acanthocotyle urolophi*; *Acanthocotyle verrilli*; *Acanthocotyle williamsi* Price, 1938. *Acanthocotyle verrilli* most closely resembles *A. atacamensis, A. gurgesiella, A. imo* and *A. urolophi*. All these species have more than 20 testes, a haptor armed with 21-39 radial rows of sclerites, and a dextral opening of the uterine pore. *Acanthocotyle urolophi* is distinguished from the other species by the form of the vitelline follicles. Unlike in *A. atacamensis, A. gurgesiella, A. imo* and *A. verrilli*, the vitelline follicles of *A. urolophi* are discrete and easy to count. The number of testes ranges from 26 to 47 (mode 36) in *A. verrilli*, 40 to 58 (mode 50) in *A. atacamensis*, 28 to 43 (mode 30) in *A. gurgesiella,* 32 to 47 (mode 41) in *A. imo* and 40 to 70 (mode 55) in *A. urolophi.* Testes of *A. verrilli* are organised in numerous rows compared to two rows in the case of *A. imo and A. gurgesiella.* Unlike in *A. atacamensis, A. imo* and *A. gurgesiella,* testes of *A. verrilli* have overlapping margins. The number of radial rows of sclerites range from 28 to 34 (mode 32) in *A. verrilli,* 28 (no variation) in *A. atacamensis,* 36 to 40 (mode 40) in *A. gurgesiella,* 30 to 35 (mode 32) in *A. imo* and 32 to 37 (mode 35) in *A. urolophi.* In specimens of *A. verrilli* possessing 28 radial rows of sclerites in the pseudohaptor, there are only 4–5 sclerites in the first row (counting from the position of the true haptor) compared to 6 in *A. atacamensis.* The difference between *A. verrilli* and *A. imo* is then visible in a smooth marginal valve of the pseudohaptor in *A. imo* compared to a distinct fringe in *A. verrilli*. *Acanthocotyle verrilli* can be distinguished from *A. gurgesiella* by the absence of an armed male genital aperture.

#### 3.1.3. Interspecific differentiation based on multivariate statistics

Principal component analysis combining metric and meristic data (see Material & Methods) was performed to examine and visualise differences between three morphologically similar species of *Acanthocotyle* for which raw data are available (Fig. 3A-C). Figure 3A shows the comparison of all three species. The first PC explained 64.5% and the second 12.3% of the variation in the dataset. In the resulting biplot, specimens of *A. verrilli* collected in this study are clearly distinguished from the other two species along the first axis and display more intraspecific variability. Figure 3B presents a PC biplot of *A. verrilli* and *A. imo* and shows clear differentiation along the first axis (PC1 explained 54.5% and PC2 17.3% of the variation in the dataset). Figure 3C presents a PCA biplot of *A. verrilli* and *A. atacamensis* and shows clear differentiation along the first axis (PC1 explained 67.1% and PC2 12.6% of the variation in the dataset). The diameter of the pseudohaptor and the total body length display the highest contribution to the separation in all datasets of all parameters.

**Fig. 3:**
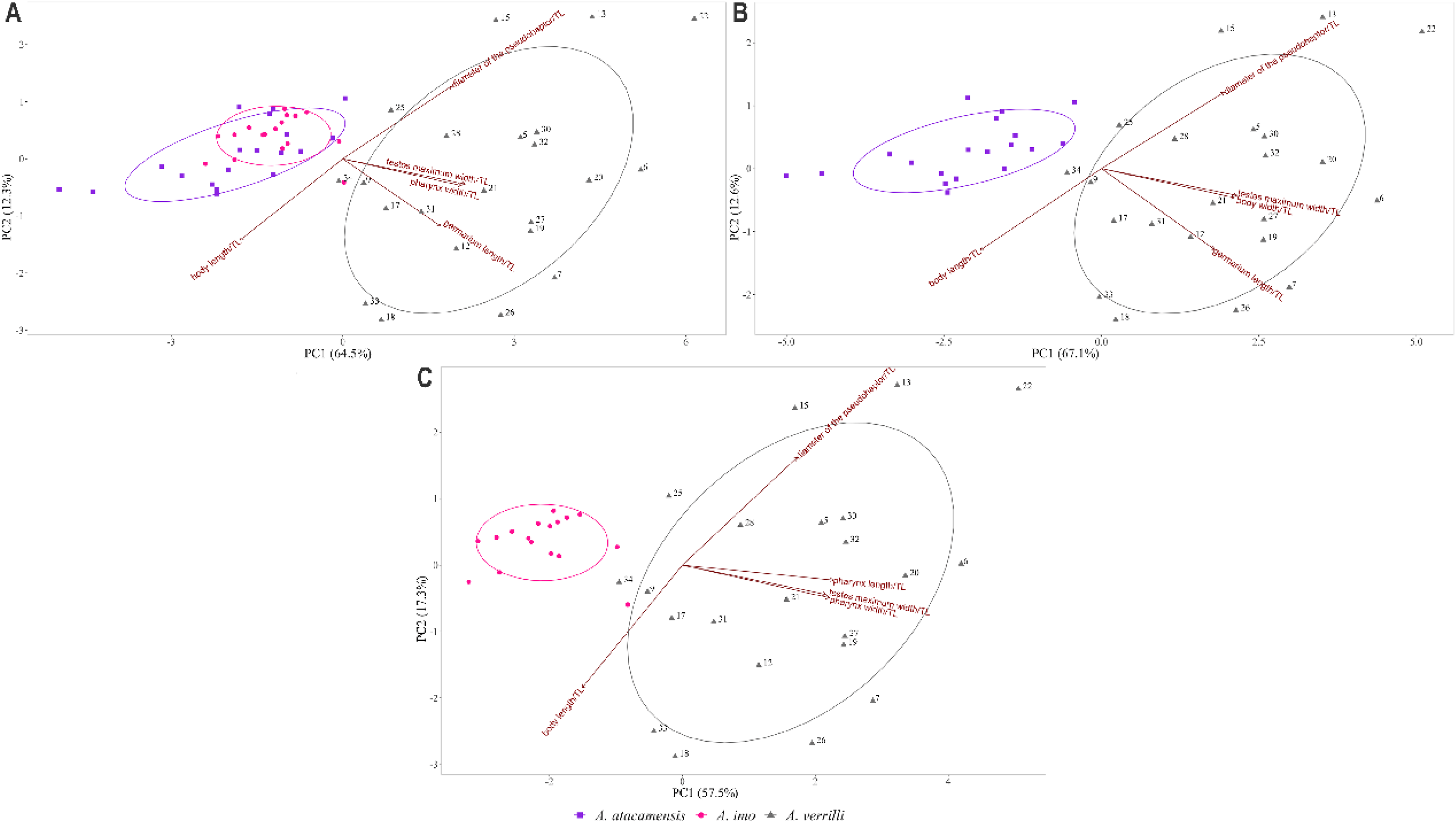
Biplots showing the interspecific differences of *Acanthocotyle* spp. based on proportional morphometric measurements standardized by the total body length. Only the first two axes are shown. A) Principal component analysis (PCA) of *A. verrilli* (this study), *A. atacamensis* [49] and *A. imo* [49]. B) PCA of *A. verrilli* (this study) and *A. atacamensis* [49]. C) PCA of *A. verrilli* (this study) and *A. imo* [49].

#### 3.1.4. Phylogenetic reconstruction

In total, three identical sequences of the 28S rDNA region from *A. verrilli* were generated in this study (Genbank accession numbers xx-xx). Phylogenetic reconstruction placed *A. verrilli* in a well-supported clade with *Acanthocotyle imo* (Fig. 4). Genetic interspecific differences between the species of *Acanthocotyle* with available 28S rDNA region sequences are presented in Table 3.

**Fig. 4:**
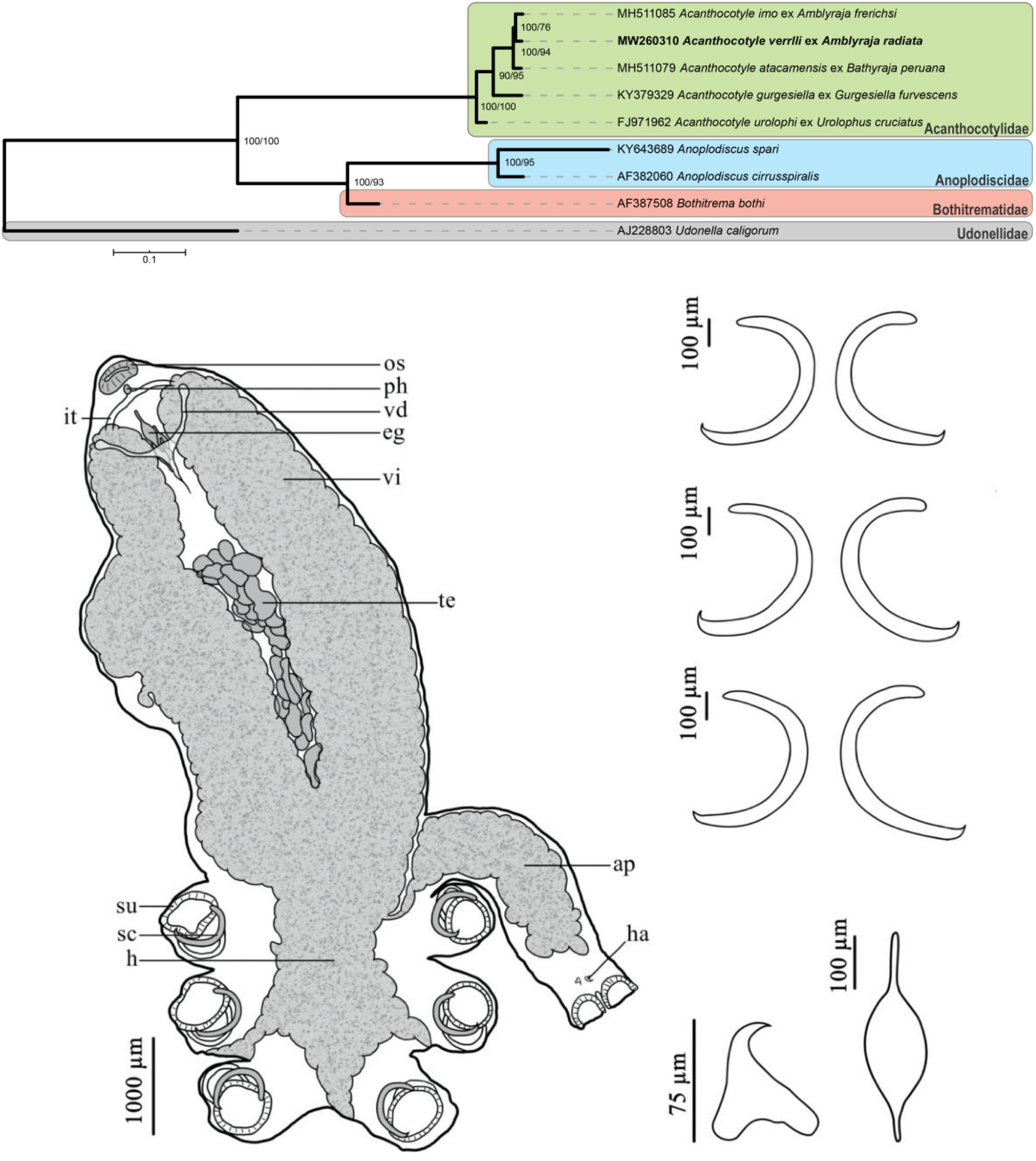
Bayesian inference phylogram based on available 28S rDNA sequences of Acanthocotylidae with specification of the host species. Representatives of three other families of Gyrodactylidea were used as outgroup. Bootstrap percentages for maximum likelihood (before slashes) and posterior probabilities for Bayesian inference (behind slashes) are shown. The scale bar indicates the expected number of substitutions per site.

**Table 3:** Uncorrected pairwise genetic distances (%) between *Acanthocotyle* spp. based on 844bp of 28S rDNA gene portion. GenBank accession numbers are given in brackets.

|  | *A. verrilli* | *A. atacamensis* | *A. gurgesiella* | *A. imo* |
| --- | --- | --- | --- | --- |
| *A. verrilli* (MW260310) |  |  |  |  |
| *A. atacamensis* (MH511079) | 0.75 |  |  |  |
| *A. gurgesiella* (KY379329) | 2.01 | 2.26 |  |  |
| *A. imo* (MH511085) | 0.25 | 0.50 | 1.76 |  |
| *A. urolophi* (FJ971962) | 3.01 | 3.01 | 3.27 | 2.76 |

### 3.2. *Rajonchocotyle emarginata* (Olsson, 1876)

Monogenea van Beneden, 1858

**Family** Hexabothriidae Price, 1942

**Genus** *Rajonchocotyle* Cerfontaine, 1899

**Type-host:** *Amblyraja radiata* (Donovan, 1808) (Chondrichtyes, Rajidae)

**Other hosts:** *Bathyraja brachyurops* (Fowler, 1910); *Bathyraja magellanica* (Philippi, 1902) (Chondrichtyes, Arhynchobatidae); *Leucoraja naevus* (Müller & Henle, 1841); *Raja brachyura* Lafont, 1871; *Raja clavata* Linnaeus, 1758; *Raja microocellata* Montagu, 1818; *Raja montagui* Fowler, 1910; *Raja undulata* Lacepède, 1802 (Chondrichtyes, Rajidae) and *Psammobatis scobina* (Philippi, 1857) (Chondrichtyes, Arhynchobatidae)

**Site on host:** Gills.

**Prevalence:** 27,3% (3 out of 11 specimens infected)

**Intensity of infection:** 3 (1-6)

**Abundance:** 0,8 (0-6).

**Type-locality:** Bohuslän Coast, Sweden

**Other localities:** Adventfjorden, Spitsbergen; Cardigan Bay, Wales; Chaleur Bay, Canada; Galway Bay, Ireland; Marine Bahusiae, Scandinavia; Mediterranean Sea, Italy; off Plymouth, UK; off Roscoff, France; Northwest coast of Spain; Patagonian Shelf; off Tromsø, Norway

**ZooBank registration**: The LSID for *Rajonchocotyle emarginata* is urn:lsid:zoobank.org:act:865F76DA-FADB-49F1-9B43-24F4AAC88256.

#### 3.2.1 Remarks

In total, 21 morphological characters including soft body parts as well as sclerotised structures were measured (see Table 4, Figs. 5&6). Body elongate with tegument covered by numerous transverse ridges organised in radial rows (Fig. 6A). Haptor symmetrical with six suckers, armed with three pairs of C-shaped haptoral sucker sclerites of similar shape and size (Fig. 5) and with a sharp hook (Fig. 6F). Peduncles of suckers of similar size. Each of the suckers contains a large sclerite ending in a hook pointing to the deep lumen (Fig. 6B&F). Sclerites form a bulge structure visible at the terminal region of each sucker (Fig. 6G). The sucker margin surmounted by a rim supporting the sclerite (Fig. 6F&H). Marginal haptoral appendix with a pair of terminal suckers with three valves (Fig. 6D) and of V-shaped hamuli possessing a sharply pointed and curved tip (Fig. 6E) situated near the distal end of appendix. Mouth subterminal, situated on the ventral side of the body and formed by the oral sucker (Fig. 6C). Pharynx spherical, reaching the posterior end of oral sucker. Intestinal tract bifurcation at the level of pharynx. Testes occupy area in the central part of the body, irregular in size and shape, number of testes not ascertainable. Other parts of male reproductive system not distinguishable. Slightly lobed ovary (paratype USNM 1337399) and Y-shaped structure of vaginal ducts. Vitellaria extending from the level of intestinal bifurcation (paratype USNM 1337399) to the posterior end of the body into the haptor. Eggs fusiform with two incipient polar filaments (Fig. 5), located at level of anterior part of vitellarium.

**Table 4:** Meristic and morphometric data for *Rajonchocotyle emarginata* ex *Amblyraja radiata* from Adventfjorden, Spitsbergen. Measurements are given in micrometers.

| *Rajonchocotyle emarginata* (n=5) |  |
| --- | --- |
| Total body length | 8497–10556 (9449)<sup>a</sup> |
| Body proper length | 6465–7943 (7093)<sup>b</sup> |
| Body width at level of ovary | 1604–2601 (2073)<sup>b</sup> |
| Anterior sucker |  |
| Length | 101–530 (280)<sup>a</sup> |
| Width | 120–858 (506)<sup>a</sup> |
| Pharynx |  |
| Length | 113–116 (115)<sup>c</sup> |
| Width | 121–121 (121)<sup>c</sup> |
| Haptor |  |
| Length | 1547–2642 (2199)<sup>a</sup> |
| Width | 2113–3337 (2556)<sup>a</sup> |
| Haptoral sclerites |  |
| Anterior sclerite length | 417–495 (455)<sup>a</sup> |
| Median sclerite length | 402–503 (447)<sup>a</sup> |
| Posterior sclerite length | 316–523 (420)<sup>a</sup> |
| Appendix |  |
| Length | 1913–4078 (3247)<sup>a</sup> |
| Width | 605–939 (749)<sup>a</sup> |
| Hamulus |  |
| Outer root length | 42–51 (46)<sup>c</sup> |
| Inner root length | 37–43 (40)<sup>c</sup> |
| Testes |  |
| Testicular field length | 1822–4075 (3088)<sup>a</sup> |
| Number | 10–36 (mode=24)<sup>a</sup> |
| Egg |  |
| Proper length | 248–303 (280)<sup>a</sup> |
| Number of filaments | 2<sup>d</sup> |
| Length of filament | 105–143 (123)<sup>c</sup> |

Notes: <sup>a</sup>(n=4); <sup>b</sup>(n=5); <sup>c</sup>(n=2); <sup>d</sup>(n=3)

**Fig. 5:**
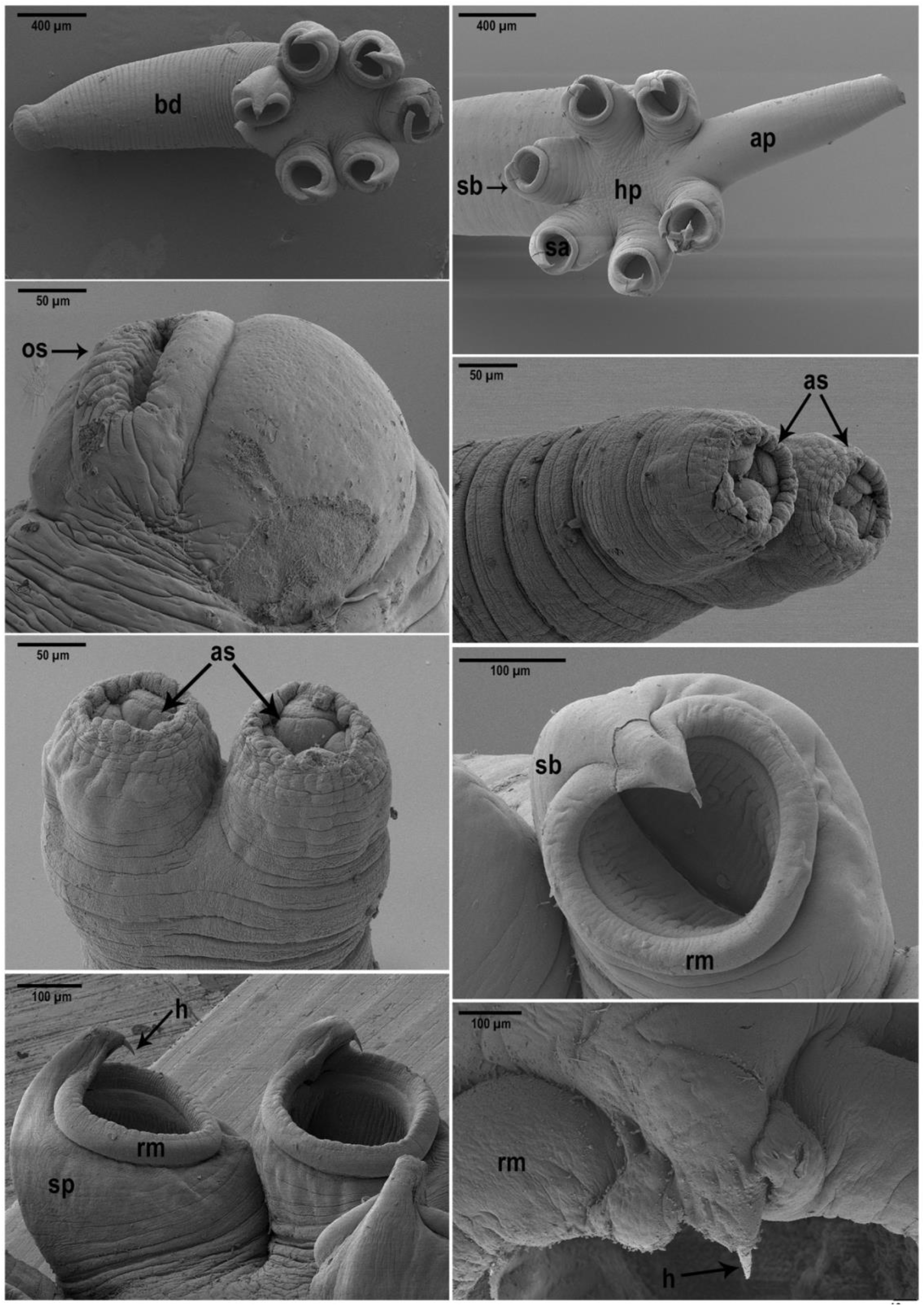
*Rajonchocotyle emarginata* ex *Amblyraja radiata*. A) Ventral view of the whole body. B) Anterior haptoral sucker sclerites. C) Middle haptoral sucker sclerites. D) Posterior haptoral sucker sclerites. E) Hamuli. F) Egg. Abbreviations: ap, appendix; eg, egg; h, haptor; ha, hamuli; it, intestine; os, oral sucker; ph, pharynx; sc, sclerite; su, sucker; te, testes; vd, vaginal duct; vi, vitellarium.

**Fig. 6:** SEM observations of *Rajonchocotyle emarginata.* A) Ventral view of the whole body, B) Haptoral structures formed by six suckers each armed with a sclerite, and appendix, C) View of the anterior part of the body with the oral sucker, D) Lateral view of the posterior part of the appendix with two terminal suckers. E) Bottom-up view on the posterior part of the appendix with two terminal suckers. F) Haptoral sucker rounded by the rim with a sclerite bulge ending in a hook. G) Side view of haptoral suckers with peduncle structure on the left side. H) Detailed view of the sucker rim and hook. ap – appendix, as – appendicular sucker, h – hook, hp – haptor, rm – sucker rim, sb – sucker bulge, sp – sucker peduncle os – oral sucker.

#### 3.2.2 Differential diagnosis

Up to now, there have been 17 genera of hexabothriid monogeneans described: *Branchotenthes* Bullard et Dippenaar, 2003; *Callorhynchocotyle* Suriano & Incorvaia, 1982; *Dasyonchocotyle* Hargis, 1955; *Epicotyle* Euzet & Maillard, 1974; *Erpocotyle* Van Beneden & Hesse, 1863; *Heteronchocotyle* Brooks, 1934; *Hexabothrium* von Nordmann, 1840; *Hypanocotyle* Chero, Cruces, Sáez, Camargo, Santos & Luque, 2018; *Mobulicola* Patella & Bullard, 2013; *Neonchocotyle* Ktari & Maillard, 1972; *Paraheteronchocotyle* Mayes, Brooks & Thorson, 1981; *Pristonchocotyle* Watson & Thorson, 1976; *Protocotyle* Euzet & Maillard, 1974; *Pseudohexabothrium* Brinkmann, 1952; *Rajonchocotyle* Cerfontaine, 1899; *Rhinobatonchocotyle* Doran, 1953 and *Squalonchocotyle* Cerfontaine, 1899. Species of *Rajonchocotyle* can be distinguished by the presence of a symmetrical haptor in comparison to *Callorhynchocotyle, Epicotyle, Heteronchocotyle, Neonchocotyle*, *Paraheteronchocotyle*, *Pristonchocotyle*, *Pseudohexabothrium* and *Rhinobatonchocotyle.* Species of *Rajonchocotyle* also differ from those within *Dasyonchocotyle* and *Hexabothrium* by having an unarmed male copulatory organ. Unlike representatives of *Branchotenthes*, *Erpocotyle*, *Hypanocotyle*, *Mobulicola* and *Squalonchocotyle* where the vagina is differentiated into muscular and glandular portions and possesses parallel vaginal ducts, species of *Rajonchocotyle* have an undifferentiated vagina and Y-shaped vaginal ducts. Species of *Rajonchocotyle* closely resemble representatives of *Protocotyle* but they differ by having undifferentiated vaginal ducts that are Y-shaped while the vaginal ducts of *Protocotyle* are also undifferentiated but parallel [51]. According to Boeger & Kritsky (1989), four species of *Rajonchocotyle* are currently considered valid: *Rajonchocotyle batis* Cerfontaine, 1899, *R. emarginata* (Olsson, 1876), *Rajonchocotyle laevis* Price, 1942 and *Rajonchocotyle wehri* Price, 1942.

*Rajonchocotyle emarginata* can be distinguished from its congeners by the total and proportional size of the haptoral sclerites. Unlike in *R. emarginata*, the size of sclerites of *R. batis* and *R. laevis* is not equal. While the anterior pair of *R. batis* is bigger than the posterior pair, sometimes even twice of the size, the posterior pair of *R. laevis* is just a bit smaller compared to the anterior pair. The difference between *R. emarginata* and *R. batis* is also visible in the shape of the anchor roots. The different pairs of haptoral sclerites of *R. wehri* and *R. emarginata* are all of equal size but the overall size of sclerites of *R. emarginata* is smaller than in *R. wehri* (median sclerites of 404–561 μm compared to 924–956 μm in *R. wehri*). *Rajonchocotyle emarginata* can also be distinguished from all its other congeners by having eggs with 2 long filaments (total egg length 353–446 μm) in comparison to the lack of polar filaments in *R. batis*, a small knob at each pole in *R. laevis* and two very short, fusiform egg filaments in *R. wehri* (total egg length 285–300 μm) [52–54].

## 4. Discussion

Invertebrate diversity is understudied in polar regions, with available information biased towards a few taxa [18,19]. In general, knowledge about the parasite fauna in cold areas remains poor, and zoonotic parasitosis received most attention so far [21]. Despite an intensified effort, reflected by the numerous parasitological surveys conducted recently [55–58], fish flatworms are rarely studied in the Svalbard archipelago. Rokicka (2009) does not mention a single monogenean infection for over 94 examined fish specimens belonging to 4 species [59]. Our study is the first record of monogenean species off Spitsbergen Island and the Svalbard archipelago.

### 4.1. Species richness and geographic distribution of *Acanthocotyle* and *Rajonchocotyle*

In total, 12 currently valid species of Acanthocotylidae have been reported out of 15 species accounting for 5.6% of the species diversity of Rajiformes. Two teleost fish species, *Reinhardtius hippoglossoides* (Walbaum, 1792) and *Sebastes alutus* (Gilbert, 1890), were also recorded as hosts for *Acanthocotyle williamsi* [60,61]. The recorded infection of *Acanthocotyle* sp. on *Narcine maculata* (Torpediniformes) is rather considered to result from transfer during fish capture [62]. With an estimation of over 800 species, the number of potential elasmobranch and holocephalan hosts of species of Acanthocotylidae is high. From less than 10% of them [63,64], only 59 species of Hexabothriidae have been described [51]. Considering the overall high species richness and rather strict host-specificity of monogeneans, the known diversity of both examined monogenean groups infecting cartilaginous fishes can be assumed to be proportionally minimal.

The overall worldwide occurrence of *Acanthocotyle* spp. seems to follow the geographical distribution of their skate hosts as summarised in Ñacari et al. [49]. Host-specificity ranges from one (9 species of *Acanthocotyle)* to five host species in *A. lobianchi.* However, the known distribution of *A. lobianchi* is currently mostly restricted to Plymouth, UK [65] with one record from Naples, Italy [66]. A similar host (four rajid species) and geographical range was reported for *A. pacifica* [67–69]. The distribution of *A. verrilli* overlaps with the cross-Atlantic occurrence of *A. radiata*. This monogenean species was further reported from two other rajid species so far (see Table 5). However, reported differences in host-specificity and distribution patterns of *Acanthocotyle* spp. are suggested to result from biased sampling toward a few host species.

**Table 5:**
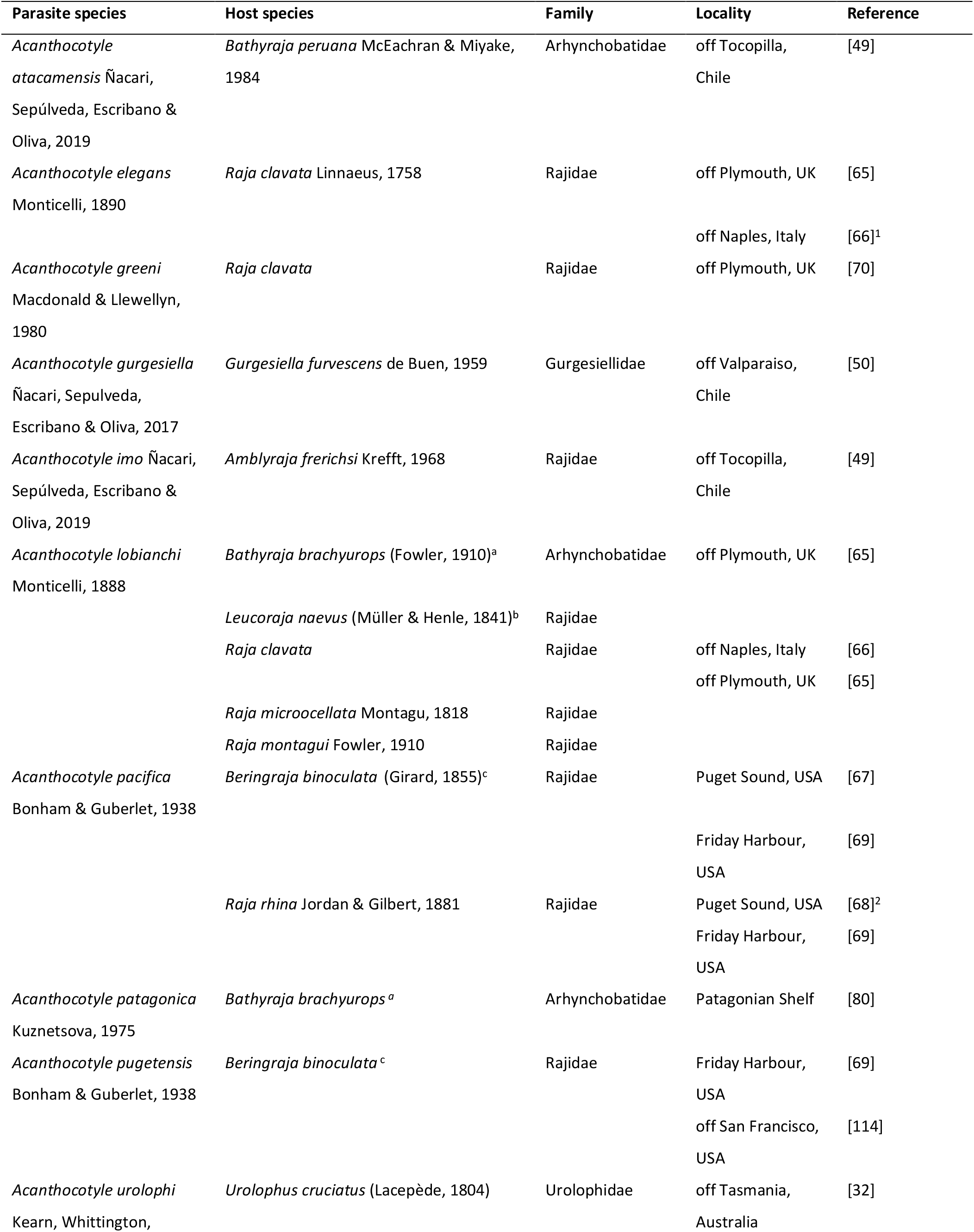
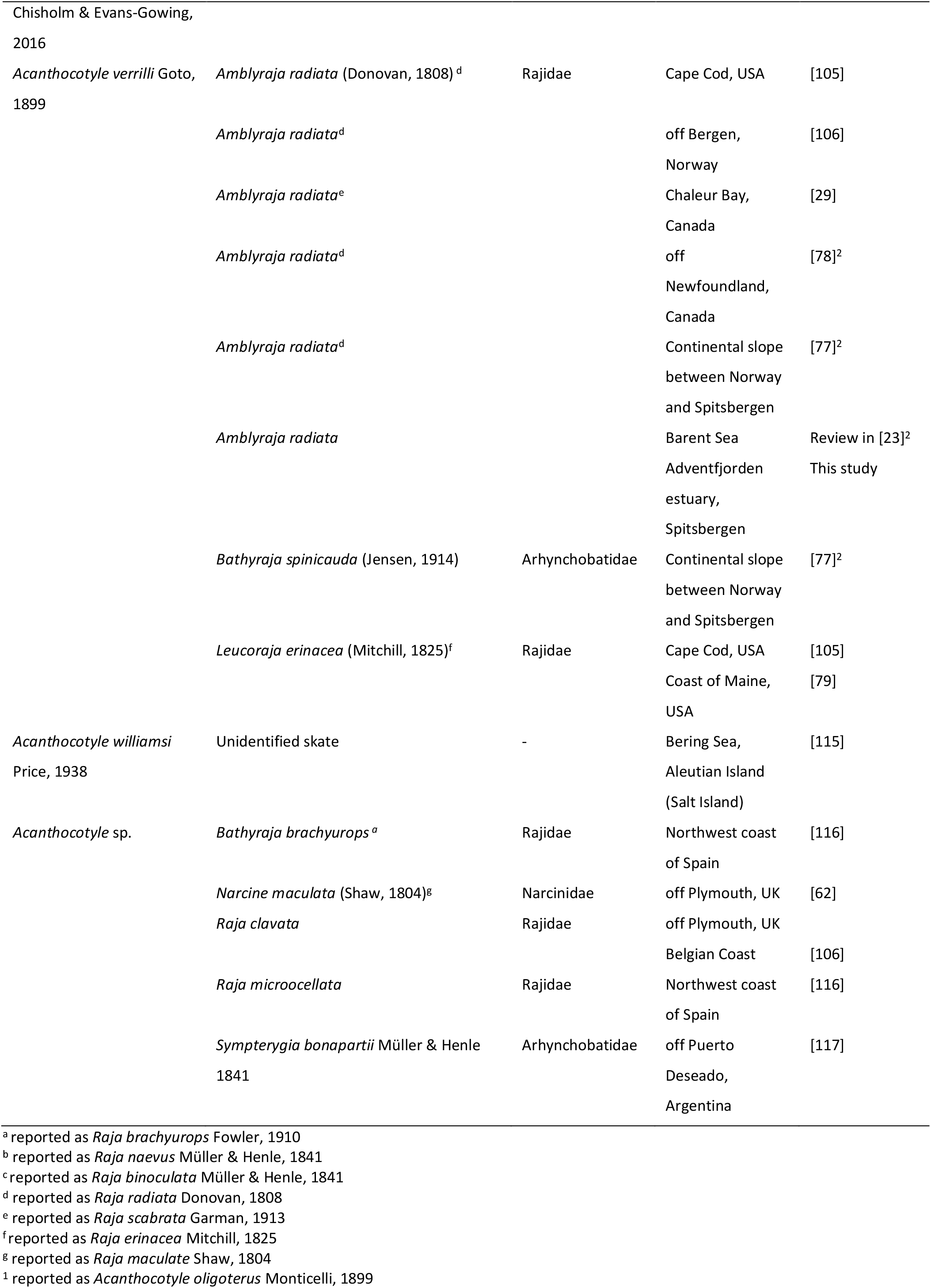
List of *Acanthocotyle* spp. with host species designation and locality of the report.

Unlike in *Acanthocotyle*, species of *Rajonchocotyle* seem to be less host specific as several representatives were recorded from different rajid hosts with a maximum number of 10 in the case of *R. emarginata.* This difference in host-specificity possibly can be driven by the site of infection (skin in *Acanthocotyle* spp. versus gills in *Rajonchocotyle* spp.) or mode of reproduction (eggs being attached to the parasite body by stalks in *Acanthocotyle* spp. [70,71] and floating eggs and free-swimming larvae in *Rajonchocotyle* spp. [72,73]. However, discovery of cryptic species that are more host specific than the species they were originally assigned to, has already changed views on parasite species richness and host-specificity [74]. Given the lack of genetic data on species of *Rajonchocotyle* and hexabothriids in general, and their close interspecific morphological similarity, the presence of cryptic species cannot be excluded. Further research is needed to verify the level of host-specificity and overall distribution patterns of both monogenean families as large parts of the host distribution remains devoid of parasitological investigation.

### 4.2. Morphological and genetic diversity

The specimens of *A. verrilli* analysed in this study did not differ in most morphological characteristics from earlier reports but some differences were observed. Goto (1899) and Sproston (1946) counted 30 or 32 radials rows of sclerites in the haptor while in our specimens the rows ranged from 28 to 34 (32 most frequently, in 14 of the 34 specimens). Thus, intraspecific variation in the number of radial rows of sclerites was reported in *A. verrilli,* as in other congeners [32,49,50]. Interestingly, a relationship between age and the number of radial rows of polyopisthocotylean monogenean species was suggested [75]. However, such pattern does not emerge for our data (see Supplementary Table S1). Moreover, because the number of rows in the pseudohaptor and the number of rows of testes can overlap, the difference between *A. verrilli* and *A. atacamensis* was revised. We propose the number of sclerites in the first row (counting from the position of the true haptor) as an additional diagnostic character. This study confirms that the morphology and size of sclerotised structures are of a high diagnostic significance in *Acanthocotyle*. Close morphological similarities of *A. verrilli* with *A. imo* and *A. atacamensis* are reflected in the genetic distance matrix (Table 3). Interestingly, the two species of *Acanthocotyle* collected from representatives of *Amblyraja* formed a clade in the phylogenetic reconstruction (Fig. 6). However, molecular data on the remaining *Acanthocotyle* spp. as well as haplotypes of *A. verrilli* from other rajid genera are needed to shed light on the evolutionary history of this parasite-host system.

As pointed out in previous studies, boundaries between hexabothriid monogenean species are mostly defined by variable characters, as they are unstable across different fixation and staining methods [51,76]. Vaughan and Christison (2012), using multivariate statistics, combined measurements of the hamulus and sucker sclerites to distinguish species of *Callorhynchocotyle.* Our study confirmed that the morphology of the hamulus and the size of sucker sclerites is of a high diagnostic significance. As a result, the combination of the proportional size of sclerotised structures and egg filaments is proposed for species identification of *Rajonchotyle* spp. Moreover, multiple staining methods should be used for correct assignment of hexabothriid monogeneans in general and species of *Rajonchocotyle* in particular.

### 4.3. Parasites of *A. radiata* as a tag for host population structure

To date, 28 helminth parasite species were reported from *A. radiata* worldwide. Even though our study was restricted to monogeneans, ongoing investigations suggest the presence of at least 9 endoparasitic helminth species infecting *A. radiata* in Svalbard (unpublished).

*Acanthocotyle verrilli* was further reported from *A. radiata* by Sproston, 1946 off Bergen, Norway, North-eastern Norwegian Sea (Rokicki and Berland, 2009), on the opposite side of the Atlantic Ocean off Newfoundland and Chaleur Bay in Canada [29,78] and the northern East coast of the USA [79]. Unlike in *A. verrilli,* the known occurrence of *R. emarginata* on *A. radiata* also spans the southern hemisphere, on the Patagonian Shelf [80]. Moreover, *R. batis* and two monogenean species from Monocotylidae were reported parasitizing on *A. radiata* in previous studies (see Table 1).

Although our results match with the previous records of monogeneans collected from *A. radiata,* the new locality off Svalbard represents the highest known latitude in the northern hemisphere those two monogenean genera have ever been recorded from (see Tables 5&6). As monogeneans display a direct life cycle and short-lived larval stage [73,81], their distribution is primarily affected by the distribution and migration patterns of their host. Given the previous reports of both monogenean species from *A. radiata* on both sides of Atlantic Ocean, historical and ongoing overseas connectivity of skate species at the Svalbard coast is proposed as suggested in Chevolot et al. [82]. Differences in life history traits such as total length and density between the populations of thorny skate were observed [5,83,84]. In contrast to the previously recorded high level of fidelity and relatively small home range (mostly fewer than 100 km [85–88]), long-term connectivity between thorny skate populations from the eastern and western part of the Atlantic facilitated by historical population expansion was recently documented [82]. These recently suggested large migratory capacities of thorny skates concur with the occasional records at depths down to 1000 m [89] with continental shelves considered as important migration barrier [16]. However, there are low levels of migration between the North Sea population and other European areas [82]. Given their expected higher mutation rate and reproduction coefficient compared to their hosts, parasitic flatworms including monogeneans were proposed as tags for historical and ongoing host migration [90–92]. However, both reported species of monogeneans infecting thorny skate are not strictly host specific (see Table 5&6). Other skate species can therefore contribute to the worldwide occurrence of *R. emarginata* and the occurrence throughout the northern hemisphere of *A. verrilli.* In general, monogeneans tend to be less host specific in pelagic and deepwater areas in comparison to littoral habitats due to the lower host availability connected also with fish population size [93–96]. On the other hand, an influence of light intensity on the larval hatching of *R. emarginata* as a result of adaptation to the behavioural differences between the hosts was proposed [72]. Such an adaptation might therefore have resulted in depth-dependent host specificity of this parasite species. Keeping in mind the rather plastic nature of currently used morphological characters in both examined monogenean families, phenotypic evaluation of any differentiation would need to be employed over a large number of specimens per population.

**Table 6:** List of *Rajonchocotyle* spp. with host species designation and locality of the report.

| Parasite species | Host species | Family | Locality | Reference |
| --- | --- | --- | --- | --- |
| *Rajonchocotyle batis*<br>Cerfontaine, 1899 | *Amblyraja hyperborea*<br>(Collett, 1879)<sup>a</sup> | Rajidae | Continental slope<br>between Norway<br>and Spitsbergen | [77] |
|  | *Amblyraja radiata*<br>(Donovan, 1808)<sup>b</sup> | Rajidae | Continental slope<br>between Norway<br>and Spitsbergen |  |
|  | *Beringraja*<br>*binoculata* (Girard, 1855)<sup>c</sup> | Rajidae | Friday Harbor,<br>Salish Sea, USA | [69] |
|  | *Bathyraja*<br>*spinicauda* (Jensen, 1914) | Arhynchobatidae | Continental slope<br>between Norway<br>and Spitsbergen | [77] |
|  | *Dipturus batis* (Linnaeus,<br>1758)<sup>d</sup> | Rajidae | off Skagerrak,<br>Denmark | [54] |
|  |  |  | off Liège, Belgium | [52] |
|  |  |  | off Ostend, Belgium |  |
|  |  |  | Porcupine Bank,<br>Ireland | [118] |
|  |  |  | off Plymouth, UK | [65] |
|  | *Raja clavata* Linnaeus, 1758 | Rajidae | off Roscoff, France | [52] |
| *Rajonchocotyle emarginata* (Olsson,<br>1876) | *Amblyraja radiata*<sup>e</sup> | Rajidae | Chaleur Bay,<br>Canada | [29] |
|  | *Amblyraja radiata*<sup>b</sup> |  | Patagonian Shelf | [111] |
|  | *Amblyraja radiata* |  | off Tromsø, Norway | [30] |
|  |  |  | Barent Sea | Review in<br>[23] |
|  |  |  | Adventfjorden<br>estuary,<br>Spitsbergen | This study |
|  | *Bathyraja brachyurops*<br>(Fowler, 1910)<sup>f</sup> | Arhynchobatidae | Patagonian Shelf | [111] |
|  | *Bathyraja magellanica*<br>(Philippi, 1902)<sup>g</sup> |  |  |  |
|  | *Leucoraja naevus* (Müller &<br>Henle, 1841)<sup>h</sup> | Rajidae | off Plymouth, UK | [65] |
|  | *Psammobatis scobina*<br>(Philippi, 1857)<sup>i</sup> | Arhynchobatidae | Patagonian Shelf | [111] |
|  | *Raja brachyura* Lafont,<br>1871 | Rajidae | off Plymouth, UK<br>Northwest coast of<br>Spain | [65]<br>[116] |
|  | *Raja clavata* | Rajidae | off Plymouth, UK<br>Mari Bahusiae,<br>Scandinavia<br>Mediterranean Sea,<br>Italy<br>Cardigan Bay, Wales<br>off Roscoff, France<br>Patagonian Shelf<br>Galway Bay, Ireland | [119] [65]<br>[120]<br>[121,122]<br>[119]<br>[53]<br>[111]<br>[123] |
|  | *Raja microocellata*<br>Montagu, 1818 | Rajidae | off Plymouth, UK<br>Northwest coast of<br>Spain | [65]<br>[116] |
|  | *Raja montagui* Fowler,<br>1910 | Rajidae | off Plymouth, UK | [65] |
|  | *Raja* sp. | Rajidae | Northwest coast of<br>Spain | [116] |
|  | *Raja undulata* Lacepède,<br>1802 | Rajidae | Northwest coast of<br>Spain |  |
| *Rajonchocotyle laevis*<br>Price, 1942 | *Dipturus laevis* (Mitchill,<br>1818)<sup>j</sup> | Rajidae | Woods Hole, USA | [53] |
| *Rajonchocotyle wehri*<br>Price, 1942 | *Raja asterias* (Delaroche,<br>1809)<sup>k</sup> | Rajidae | Friday Harbor, USA | [53] |

<sup>a</sup> reported as *Raja hyperborea* Collett, 1879
<sup>b</sup> reported as *Raja radiata* Donovan, 1808
<sup>c</sup> reported as *Raja binoculata* Girard, 1855
<sup>d</sup> reported as *Raja batis* Linnaeus, 1758
<sup>e</sup> reported as *Raja scabrata* Garman, 1913
<sup>f</sup> reported as *Raja brachyurops* Fowler, 1910
<sup>g</sup> reported as *Raja magellanica* Philippi, 1902
<sup>h</sup> reported as *Raja naevus* Müller & Henle
<sup>i</sup> reported as *Raja scobina* Philippi, 1857
<sup>j</sup> reported as *Raja laevis* Mitchill, 1818
<sup>k</sup> reported as *Raja stellata* Delaroche, 1809

More variable genetic markers would need to be applied in order to investigate the historical connectivity of the parasite populations in Svalbard and other areas and evaluate the level of geographically/host species driven differentiation.

## Acknowledgement

We would like to thank Marek Brož, Alena Sucháčková, Martins Briedis and Eva Myšková for the help with fish collection and hospitability of all the crew members at the Czech Nostoc field station.

## Supplementary material

Table S1: Raw morphometric data for *Acanthocotyle verrilli* ex *Amblyraja radiata.* Measurement are given in micrometers.

Table S2: Raw morphometric data for *Rajonchocotyle emarginata* ex *Amblyraja radiata*.

Measurement are given in micrometers.

## References

[1] C. Sguotti, C.P. Lynam, B. García-Carreras, J.R. Ellis, G.H. Engelhard, Distribution of skates and sharks in the North Sea: 112 years of change, Glob. Chang. Biol. 22 (2016) 2729–2743. https://doi.org/10.1111/gcb.13316.

[2] D.P. Packer, C.A. Zetlin, J.J. Vitaliano, Thorny skate, *Amblyraja radiata,* life history and habitat characteristics, 2003.

[3] K. Brander, Disappearance of common skate *Raia batis* from Irish Sea, Nature. 290 (1981) 48–49. https://doi.org/10.1038/290048a0.

[4] E.J. Heist, A review of population genetics in sharks, in: Am. Fish. Soc. Symp., 1999: pp. 161–168.

[5] J. Sulikowski, J. Kneebone, S. Elzey, J. Jurek, P. Danley, W. Howell, P. Tsang, Age and growth estimates of the thorny skate *(Amblyraja radiata)* in the western Gulf of Maine, Fish. Bull. (2005).

[6] P.A. Walker, G. Howlett, R. Millner, Distribution, movement and stock structure of three ray species in the North Sea and eastern English Channel, ICES J. Mar. Sci. 54 (1997) 797–808. https://doi.org/10.1006/jmsc.1997.0223.

[7] H.B. Bigelow, W.C. Schroeder, Sawfishes, Guitarfishes, Skates and Rays, Chimaeroids: Part 2, Yale University Press, 2018.

[8] J.R. Ellis, N.K. Dulvy, S. Jennings, M. Parker-Humphreys, S.I. Rogers, Assessing the status of demersal elasmobranchs in UK waters: a review, J. Mar. Biol. Assoc. United Kingdom. 85 (2005) 1025–1047. https://doi.org/10.1017/S0025315405012099.

[9] J.R. Ellis, A. Cruz-Martínez, B.D. Rackham, S.I. Rogers, The distribution of chondrichthyan fishes around the British Isles and implications for conservation, J. Northwest Atl. Fish. Sci. 35 (2005) 195–213. https://doi.org/10.2960/j.v35.m485.

[10] D.P. Swain, H.P. Benoit, Change in habitat associations and geographic distribution of thorny skate *(Amblyraja radiata)* in the southern Gulf of St Lawrence: density-dependent habitat selection or response to environmental change?, Fish. Oceanogr. 15 (2006) 166–182. https://doi.org/10.1111/j.1365-2419.2006.00357.x.

[11] J.D. Mceachran, M.R. de Carvalho, Batoid Fishes, in: K.E. Carpenter, V.H. Niem (Eds.), Living Mar. Resour. West. Cent. Pacific, K.E. Carpenter, Rome, 1999: pp. 508–589.

[12] F. Cousteau, Ocean: the definitive visual guide, New York, 2014.

[13] M. Jakobsson, A. Grantz, Y. Kristoffersen, The Arctic Ocean: boundary conditions and background information, in: R. Stein, R.W. MacDonald (Eds.), Org. Carbon Cycle Arct. Ocean, Springer Berlin Heidelberg, Berlin, Heidelberg, 2004: pp. 1–32. https://doi.org/10.1007/978-3-642-18912-8_1.

[14] W.B. Harland, Chapter 1: Svalbard, Geol. Soc. Mem. 17 (1997) 3–15. https://doi.org/10.1144/GSL.MEM.1997.017.01.01.

[15] C. Michel, Marine ecosystems, in: H. Meltofte (Ed.), Arct. Biodivers. Assess., Aarhus, 2013: pp. 486–527. https://doi.org/10.1201/9780203757222-17.

[16] C.W. Cunningham, T.M. Collins, B. Schierwater, B. Streit, G.P. Wagner, R. DeSalle, Molecular ecology and evolution, approaches and application, Switz. Birkhauser Verlad Base. 405 (1994) 433.

[17] A. Clarke, The Polar Deep Seas, in: P.A. Tyler (Ed.), Ecosyst. Deep Ocean., Elsevier Science B.V., Amsterdam, 2003: pp. 241–262.

[18] S.J. Coulson, Terrestrial and freshwater invertebrate fauna of the high arctic archipelago of Svalbard, Zootaxa. 1448 (2007) 41–58. https://doi.org/10.11646/zootaxa.1448.1.2.

[19] R. Palerud, B. Gulliksen, T. Brattegard, J.A. Sneli, W. Vader, The marine macroorganisms in Svalbard waters. A catalogue of the terrestrial and marine animals of Svalbard., Nor. Polar Institute, Skr. 201 (2004) 5–56.

[20] K. Rohde, Ecology and biogeography of marine parasites., Adv. Mar. Biol. 43 (2002) 1–83. https://doi.org/10.1016/s0065-2881(02)43002-7.

[21] J. Dupouy-Camet, Parasites of cold climates: A danger or in danger?, Food Waterborne Parasitol. 4 (2016) 1–3. https://doi.org/10.1016/j.fawpar.2016.07.004.

[22] K. V. Galaktionov, Patterns and processes influencing helminth parasites of Arctic coastal communities during climate change, J. Helminthol. 91 (2017) 387–408. https://doi.org/10.1017/S0022149X17000232.

[23] A.B. Karasev, A catalogue of parasites of the Barents Sea fishes., Izd-vo PINRO, Murmansk, 2003.

[24] S.A. Murzina, S.G. Sokolov, S.N. Pekkoeva, E.P. Ieshko, N.N. Nemova, R. Kristoffersen, S. Falk-Petersen, First data on the parasite fauna of daubed shanny *Leptoclinus maculatus* (Fries 1838) (Actinopterygii, Perciformes: Stichaeidae) in Svalbard waters, Polar Biol. 42 (2019) 831–834. https://doi.org/10.1007/s00300-018-02448-2.

[25] J.N. Caira, C.J. Healy, Elasmobranchs as hosts of metazoan parasites, in: J.C. Carrier, J.A. Musick, M.R. Heithaus (Eds.), Biol. Sharks Their Relat., CRC Press, London, 2004: pp. 523–552. https://doi.org/10.1201/9780203491317.ch18.

[26] O.N. Pugachev, P.I. Gerasev, A.V. Gussev, R. Ergens, I. Khotenowsky, Guide to Monogenoidea of freshwater fish of Palaeartic and Amur regions, Ledizione-LediPublishing, Milan, 2009.

[27] I.D. Whittington, Diversity ‘‘down under’’: monogeneans in the Antipodes (Australia) with a prediction of monogenean biodiversity worldwide, Int J Parasitol. 28 (1998). https://doi.org/10.1016/S0020-7519(98)00064-2.

[28] N.G. Sproston, A synopsis of the monogenetic trematodes., Trans. Zool. Soc. London. 25 (1946) 185–600.

[29] A.F. Heller, Parasites of cod and other marine fish from the baie de Chaleur region, Can. J. Res. 27 (1949) 243–264. https://doi.org/10.1139/cjr49d-022.

[30] L.G. Poddubnaya, W. Hemmingsen, D.I. Gibson, Ultrastructural observations of the attachment organs of the monogenean *Rajonchocotyle emarginata* (Olsson, 1876) (Polyopisthocotylea: Hexabothriidae), a gill parasite of rays, Parasitol. Res. 115 (2016) 2285–2297. https://doi.org/10.1007/s00436-016-4973-x.

[31] R. Ergens, J. Lom, Causative agents of fish diseases, Academia, Prague, 1970.

[32] G. Kearn, I. Whittington, L. Chisholm, R. Evans-Gowing, A new species of *Acanthocotyle* Monticelli, 1888 (Platyhelminthes: Monogenea: Acanthocotylidae) from the ventral skin of the banded stingaree, *Urolophus cruciatus* (Lacépède, 1804), from Tasmania, Australia, Acta Parasitol. 61 (2016) 607–613. https://doi.org/10.1515/ap-2016-0081.

[33] S.A. Bullard, S.M. Dippenaar, *Branchotenthes robinoverstreetin* gen. and n. sp. (Monogenea: Hexabothriidae) from gill filaments of the bowmouth guitarfish, *Rhina ancylostoma* (Rhynchobatidae), in the Indian Ocean, J. Parasitol. 89 (2003) 595–601.

[34] F.A. Sepúlveda, M.T. González, M.E. Oliva, Two new species of *Encotyllabe* (Monogenea: Capsalidae) based on morphometric and molecular evidence: parasites of two inshore fish species of northern Chile, J. Parasitol. 100 (2014) 344–349. https://doi.org/10.1645/13-230.1.

[35] R Core Team, R: A language and environment for statistical computing. R Foundation for Statistical Computing, the R Foundation for Statistical Computing, Vienna, Austria, 2019. https://www.r-project.org/.

[36] H. Wickham, Ggplot2: elegant graphics for data analysis, Springer, 2009.

[37] S.-O. Fankoua, A.R. Bitja Nyom, D. ne dort Bahanak, C.F. Bilong Bilong, A. Pariselle, Influence of preservative and mounting media on the size and shape of monogenean sclerites, Parasitol. Res. 116 (2017) 2277–2281. https://doi.org/10.1007/s00436-017-5534-7.

[38] N. Hassouna, B. Michot, J.-P. Bachellerie, D. Narbonne, The complete nucleotide sequence of mouse 28S rRNA gene. Implications for the process of size increase of the large subunit rRNA in higher eukaryotes., Nucleic Acids Res. 12 (1984) 3563–3583. https://doi.org/10.1093/nar/12.8.3563.

[39] S. Kumar, G. Stecher, K. Tamura, J. Gerken, E. Pruesse, C. Quast, T. Schweer, J. Peplies, W. Ludwig, F. Glockner, MEGA7: Molecular evolutionary genetics analysis Version 7.0 for bigger datasets, Mol. Biol. Evol. 33 (2016) 1870–1874. https://doi.org/10.1093/molbev/msw054.

[40] R.C. Edgar, MUSCLE: Multiple sequence alignment with high accuracy and high throughput., Nucleic Acids Res. 32 (2004) 1792–1797. https://doi.org/10.1093/nar/gkh340.

[41] W.A. Boeger, D.C. Kritsky, Phylogenetic relationships of the Monogenoidea, Syst. Assoc. Spec. Vol. 60 (2001) 92–102.

[42] J. Castresana, Selection of conserved blocks from multiple alignments for their use in phylogenetic analysis, Mol. Biol. Evol. 17 (2000) 540–552. https://doi.org/10.1093/oxfordjournals.molbev.a026334.

[43] M. Hasegawa, H. Kishino, T. aki Yano, Dating of the human-ape splitting by a molecular clock of mitochondrial DNA, J. Mol. Evol. 22 (1985) 160–174. https://doi.org/10.1007/BF02101694.

[44] D. Darriba, G.L. Taboada, R. Doallo, D. Posada, jModelTest 2: more models, new heuristics and parallel computing, Nat. Methods. 9 (2012) 772–772. https://doi.org/10.1038/nmeth.2109.

[45] F. Ronquist, M. Teslenko, P. van der Mark, D.L. Ayres, A. Darling, S. Hohna, B. Larget, L. Liu, M.A. Suchard, J.P. Huelsenbeck, MrBayes 3.2: Efficient Bayesian phylogenetic inference and model choice across a large model space, Syst. Biol. 61 (2012) 539–542. https://doi.org/10.1093/sysbio/sys029.

[46] A. Rambaut, M.A. Suchard, A.J. Drummond, Tracer v1.6, (2014). http://beast.bio.ed.ac.uk.

[47] A. Stamatakis, RAxML version 8: A tool for phylogenetic analysis and post-analysis of large phylogenies, Bioinformatics. 30 (2014) 1312–1313. https://doi.org/10.1093/bioinformatics/btu033.

[48] ICZN, Amendment of Articles 8, 9, 10, 21 and 78 of the International Code of Zoological Nomenclature to expand and refine methods of publication, Zootaxa. (2012) 1–7. https://doi.org/10.3897/zookeys.219.3994.

[49] L.A. Ñacari, F.A. Sepúlveda, R. Escribano, M.E. Oliva, Two new species of *Acanthocotyle* Monticelli, 1888 (Monogenea: Acanthocotylidae), parasites of two deep-sea skates (Elasmobranchii: Rajiformes) in the South-East Pacific, Parasit. Vectors. 12 (2019) 512. https://doi.org/10.1186/s13071-019-3756-5.

[50] L.A. Ñacari, F.A. Sepulveda, R. Escribano, M.E. Oliva, *Acanthocotyle gurgesiella* n. sp. (Monogenea: Acanthocotylidae) from the deep-sea skate *Gurgesiella furvescens* (Rajidae) in the south-eastern Pacific, J. Helminthol. 92 (2018) 223–227. https://doi.org/10.1017/S0022149X17000220.

[51] W.A. Boeger, D.C. Kritsky, Phylogeny, coevolution, and revision of the Hexabothriidae Price, 1942 (Monogenea), Int. J. Parasitol. 19 (1989) 425–440. https://doi.org/10.1016/0020-7519(89)90099-4.

[52] P. Cerfontaine, Contribution à l’étude des Octocotylidés: Les Onchocotylinae, Arch. Biol. (Liege). 16 (1899) 345–478. https://doi.org/10.1017/CBO9781107415324.004.

[53] E.W. Price, North American monogenetic trematodes. V. The family Hexabothriidae, n. n. (Polystomatoidea), Proc. Helminthol. Soc. Wash. 9 (1942) 39–56. https://doi.org/10.1126/science.35.901.553.

[54] Olsson Peter, Bidrag till Skandinaviens helminthfauna, Sven. Vetenskaps–Akademiens Handl. 14 (1876) 1–35.

[55] E. Myšková, M. Brož, E. Fuglei, J. Kvičerová, A. Mácová, B. Sak, M. Kváč, O. Ditrich, Gastrointestinal parasites of arctic foxes *(Vulpes lagopus)* and sibling voles *(Microtus levis)* in Spitsbergen, Svalbard, Parasitol. Res. 118 (2019) 3409–3418. https://doi.org/10.1007/s00436-019-06502-8.

[56] J. Elsterová, J. Černý, J. Müllerová, R. Šíma, S.J. Coulson, E. Lorentzen, H. Strøm, L. Grubhoffer, Search for tick-borne pathogens in the Svalbard Archipelago and Jan Mayen, Polar Res. 34 (2015) 27466. https://doi.org/10.3402/polar.v34.27466.

[57] K.W. Prestrud, K. Asbakk, E. Fuglei, T. Mørk, A. Stien, E. Ropstad, M. Tryland, G.W. Gabrielsen, C. Lydersen, K.M. Kovacs, M.J.J.E. Loonen, K. Sagerup, A. Oksanen, Serosurvey for *Toxoplasma gondii* in arctic foxes and possible sources of infection in the high Arctic of Svalbard., Vet. Parasitol. 150 (2007) 6–12. https://doi.org/10.1016/j.vetpar.2007.09.006.

[58] H. Henttonen, E. Fuglei, C.N. Gower, V. Haukisalmi, R.A. Ims, J. Niemimaa, N.G. Yoccoz, *Echinococcus multilocularis* on Svalbard: Introduction of an intermediate host has enabled the local life-cycle, Parasitology. 123 (2001) 547–552. https://doi.org/10.1017/S0031182001008800.

[59] M. Rokicka, Report on species of *Gyrodactylus* Nordmann, 1832, distribution in polar regions, Polar Sci. 3 (2009) 203–206. https://doi.org/10.1016/j.polar.2009.07.001.

[60] A.D. Sekerak, H.P. Arai, Helminths of *Sebastes alutus* (Pisces: Teleostei) from the northeastern Pacific., Can. J. Zool. 51 (1973) 475–477. https://doi.org/10.1139/z73-071.

[61] J. Wierzbicka, W. Piasecki, Redescription of *Pseudacanthocotyla williamsi* (price, 1938) (Monogenea) from Greenland halibut, *Reinhardtius hippoglossoides* (Walbaum, 1792), Acta Ichthyol. Piscat. 30 (2000) 93–97. https://doi.org/10.3750/AIP2000.30.2.09.

[62] H.A. Baylis, E.I. Jones, Some records of parasitic worms from marine fishes at Plymouth, J. Mar. Biol. Assoc. United Kingdom. 18 (1933) 627–634. https://doi.org/10.1017/S0025315400043940.

[63] D.A. Ebert, L.J. V. Compagno, Biodiversity and systematics of skates (Chondrichthyes: Rajiformes: Rajoidei), Environ. Biol. Fishes. 80 (2007) 5–18. https://doi.org/10.1007/978-1-4020-9703-4_2.

[64] D.A. Ebert, K.E. van Hees, Beyond Jaws: rediscovering the ‘lost sharks’ of southern Africa, African J. Mar. Sci. 37 (2015) 141–156. https://doi.org/10.2989/1814232X.2015.1048730.

[65] J. Llewellyn, J.E. Green, G.C. Kearn, A check-list of monogenean (platyhelminth) parasites of Plymouth hosts, J. Mar. Biol. Assoc. United Kingdom. 64 (1984) 881–887. https://doi.org/10.1017/S0025315400047299.

[66] F.S. Monticelli, Il genere *Acanthocotyle*, Arch. Parasitol. 2 (1899) 75 – 120.

[67] K. Bonham, J.E. Guberlet, Ectoparasitic trematodes of puget sound fishes *Acanthocotyle*, Am. Midl. Nat. 20 (1938) 590. https://doi.org/10.2307/2420295.

[68] M. Love, M. Moser, A checklist of parasites of California, Oregon, and Washington marine and estuarine fishes, Fac. Publ. from Harold W. Manter Lab. Parasitol. (1983). https://digitalcommons.unl.edu/parasitologyfacpubs/750 (accessed May 26, 2020).

[69] E.S. Robinson, Some monogenetic trematodes from marine fishes of the Pacific, Trans. Am. Microsc. Soc. 80 (1961) 235. https://doi.org/10.2307/3223640.

[70] S. Macdonald, J. Llewellyn, Reproduction in *Acanthocotyle greeni* n. sp. (Monogenea) from the skin of *Raia* spp. at Plymouth, J. Mar. Biol. Assoc. United Kingdom. 60 (1980) 81–88. https://doi.org/10.1017/S0025315400024139.

[71] I.D. Whittington, G.C. Kearn, Effects of urea analogs on egg hatching and movement of unhatched larvae of monogenean parasite *Acanthocotyle lobianchi* from skin of *Raja montagui*, J. Chem. Ecol. 16 (1990) 3523–3529. https://doi.org/10.1007/BF00982115.

[72] I.D. Whittington, G.C. Kearn, Rhythmical hatching and oncomiracidial behaviour in the hexabothriid monogenean *Rajonchocotyle emarginata* from the gills of *Raja* spp., J. Mar. Biol. Assoc. United Kingdom. 66 (1986) 93–111. https://doi.org/10.1017/S0025315400039679.

[73] I.D. Whittington, L.A. Chisholm, K. Rohde, The larvae of Monogenea (Platyhelminthes), Adv. Parasitol. 44 (1999) 139–232. https://doi.org/10.1016/S0065-308X(08)60232-8.

[74] M.P.M. Vanhove, T. Huyse, Host specificity and species jumps in fish-parasite systems, in: Parasite Divers. Diversif. Evol. Ecol. Meets Phylogenetics, Cambridge University Press, 2015: pp. 401–419. https://doi.org/10.1017/CBO9781139794749.024.

[75] J. Lou Justine, A. Grugeaud, Does the number of sclerotised structures used for the systematics of monogeneans change with age? A study of the monocotylid *Dendromonocotylepipinna*, Parasitol. Res. 107 (2010) 1509–1514. https://doi.org/10.1007/s00436-010-2019-3.

[76] D. Vaughan, K. Christison, Towards addressing the current state of confusion within the Hexabothriidae Price, 1942 (1908): *Callorhynchocotyle* Suriano & Incorvaia, 1982 (Monogenea: Hexabothriidae) re-visited, with the preliminary evaluation of novel parameters for measuring h, Zootaxa. 34 (2012) 1–34. https://doi.org/10.11646/zootaxa.3229.1.1.

[77] J. Rokicki, B. Berland, Some helminth and copepod parasites of three rajid species from the continental slope of the north-eastern Norwegian Sea, Acta Parasitol. 46 (2009) 12–17.

[78] W. Threlfall, Some parasites from elasmobranchs in Newfoundland, J. Fish. Res. Board Canada. 26 (1969) 805–811. https://doi.org/10.1139/f69-078.

[79] H.W. Manter, Some North American fish trematodes, Illinois Biol. Monogr. 10 (1926) 1–138.

[80] E.G. Kuznetsova, The monogenetic trematodes of cartilaginous fish of the Patagonian Shelf of the Atlantic Ocean, Tr. Uprk Kadrov i Uchenykh Zaved. Minist. Rybn. Khozyaistva SSR. 26 (1971) 12–21.

[81] G.C. Kearn, The life-cycles and larval development of some acanthocotylids (Monogenea) from Plymouth rays, Parasitology. 57 (1967) 157–167. https://doi.org/10.1017/S0031182000071961.

[82] M. Chevolot, P.H.J. Wolfs, J. Pálsson, A.D. Rijnsdorp, W.T. Stam, J.L. Olsen, Population structure and historical demography of the thorny skate *(Amblyraja radiata,* Rajidae) in the North Atlantic, Mar. Biol. 151 (2007) 1275–1286. https://doi.org/10.1007/s00227-006-0556-1.

[83] J. Sulikowski, J. Kneebone, S. Elzey, J. Jurek, P. Danley, W. Howell, P. Tsang, The reproductive cycle of the thorny skate *(Amblyraja radiata)* in the western Gulf of Maine, Fish. Bull. (2005).

[84] W. Templeman, Development & occurrence and characteristics of egg capsules of the thorny skate & *Raja radiata* in the Northwest Atlantic, 1982.

[85] N. Daan, H.J.L. Heessen, R. ter Hofstede, North Sea Elasmobranchs: distribution, abundance and biodiversity, (2005).

[86] N.K. Dulvy, J.D. Metcalfe, J. Glanville, M.G. Pawson, J.D. Reynolds, Fishery stability, local extinctions, and shifts in community structure in skates, Conserv. Biol. 14 (2000) 283–293. https://doi.org/10.1046/j.1523-1739.2000.98540.x.

[87] W. Templeman, Migrations of thorny skate, *Raja radiata,* tagged in the Newfoundland area, J. Northw. Atl. Fish. ScL. 5 (1984) 55–63.

[88] P. Walker, G. Howlett, R. Millner, Distribution, movement and stock structure of three ray species in the North Sea and eastern English Channel, ICES J. Mar. Sci. 54 (1997) 797–808. https://doi.org/10.1006/jmsc.1997.0223.

[89] M. Stehmann, D. Bürkel, Rajidae, in: P. Whitehead, M. Bauchot, J.-C. Hureau, J. Nielsen, E. Tortonese (Eds.), Fishes North-Eastern Atl. Mediterr. Vol. I, 1994: pp. 163–196.

[90] C. Nieberding, S. Morand, R. Libois, J.R. Michaux, A parasite reveals cryptic phylogeographic history of its host., Proceedings. Biol. Sci. 271 (2004) 2559–68. https://doi.org/10.1098/rspb.2004.2930.

[91] T. Huyse, R. Poulin, A. Théron, Speciation in parasites: a population genetics approach, Trends Parasitol. 21 (2005) 469–475. https://doi.org/10.1016/j.pt.2005.08.009.

[92] M. Barson, I. Přikrylová, M.P.M. Vanhove, T. Huyse, Parasite hybridization in African *Macrogyrodactylus* spp. (Monogenea, Platyhelminthes) signals historical host distribution., Parasitology. 137 (2010) 1585–1595. https://doi.org/10.1017/S0031182010000302.

[93] M. Bueno-Silva, W. a. Boeger, M.R. Pie, Choice matters: Incipient speciation in *Gyrodactylus corydori* (Monogenoidea: Gyrodactylidae), Int. J. Parasitol. 41 (2011) 657–667. https://doi.org/10.1016/j.ijpara.2011.01.002.

[94] N. Kmentová, M. Gelnar, M. Mendlová, M. Van Steenberge, S. Koblmüller, M.P.M. Vanhove, Reduced host-specificity in a parasite infecting non-littoral Lake Tanganyika cichlids evidenced by intraspecific morphological and genetic diversity, Sci. Rep. 6 (2016) 39605. https://doi.org/10.1038/srep39605.

[95] N. Kmentová, M. Van Steenberge, J.A.R. Raeymaekers, S. Koblmüller, P.I. Hablützel, F. Muterezi Bukinga, T. Mulimbwa N’sibula, P. Masilya Mulungula, B. Nzigidahera, G. Ntakimazi, M. Gelnar, M.P.M. Vanhove, Monogenean parasites of sardines in Lake Tanganyika: diversity, origin and intra-specific variability, Contrib. to Zool. 87 (2018) 105–132.

[96] S. Morand, B.R. Krasnov, The biogeography of host-parasite interactions, Oxford University Press, 2010.

[97] L.A. Chisholm, T.J. Hansknecht, I.D. Whittington, R.M. Overstreet, A revision of the Calicotylinae Monticelli, 1903 (Monogenea: Monocotylidae), Syst. Parasitol. 38 (1997) 159–183. https://doi.org/10.1023/A:1005844306178.

[98] H.S. Randhawa, G.W. Saunders, M.D.B. Burt, Establishment of the onset of host specificity in four phyllobothriid tapeworm species (Cestoda: Tetraphyllidea) using a molecular approach, Parasitology. 134 (2007) 1291–1300. https://doi.org/10.1017/S0031182007002521.

[99] H.S. Randhawa, M.D.B. Burt, Determinants of host specificity and comments on attachment site specificity of tetraphyllidean cestodes infecting rajid skates from the Northwest Atlantic, J. Parasitol. 94 (2008) 436–461. https://doi.org/10.1645/ge-1180.1.

[100] S.S. Hendrix, Marine flora and fauna of the eastern United Stats Platyhelminthes: Monogenea, Fish. Bull. 121 (1994) 1–106.

[101] I. Beveridge, R.A. Campbell, Revision of the *Grillotia erinaceus* (van Beneden, 1858) species complex (Cestoda: Trypanorhyncha), with the description of *G. brayi* n. sp., Syst. Parasitol. 68 (2007) 1–31. https://doi.org/10.1007/s11230-006-9082-2.

[102] J. Rokicki, B. Berland, Some helminth and copepod parasites of three rajid species from the continental slope of the north-eastern Norwegian Sea, Acta Parasitol. 46 (2001) 12–17.

[103] H.S. Randhawa, Numerical and functional responses of intestinal helminths in three rajid skates: Evidence for competition between parasites?, Parasitology. 139 (2012) 1784–1793. https://doi.org/10.1017/S0031182012001035.

[104] D.T.J. Littlewood, K. Rohde, K.A. Clough, The phylogenetic position of *Udonella* (Platyhelminthes), Int. J. Parasitol. 28 (1998) 1241–1250. https://doi.org/10.1016/S0020-7519(98)00108-8.

[105] S. Goto, Notes on some exotic species of ectoparasitic trematodes, J. Coll. Sci. (1899) 263–295.

[106] A.J. Birkmann, Contribution to our knowledge of the monogenetic trematodes, Bergen. Mus. Aarb. Naturvitenskapligrekke. 1 (1940) 1–117.

[107] B. Dawes, I. Griffiths, The enigmatical trematode *Dictyocotyle coeliaca*, Nature. 182 (1958) 1033–1034. https://doi.org/10.1038/1821033a0.

[108] L.A. Chisholm, J.A.T. Morgan, R.D. Adlard, I.D. Whittington, Phylogenetic analysis of the monocotylidae (Monogenea) inferred from 28S rDNA sequences, Int. J. Parasitol. 31 (2001) 1537–1547. https://doi.org/10.1016/S0020-7519(01)00313-7.

[109] K. Rohde, C. Hefford, J.T. Ellis, P.R. Baverstock, A.M. Johnson, N.A. Watson, S. Dittmann, Contributions to the phylogeny of platyhelminthes based on partial sequencing of 18S ribosomal DNA, Int. J. Parasitol. 23 (1993) 705–724. https://doi.org/10.1016/0020-7519(93)90067-9.

[110] C.P. Keeling, M.D.B. Burt, *Echeneibothrium canadensis* n.sp. (Tetraphyllidea: Phyllobothriidae) in the spiral intestine of the thorny skate *(Raja radiata)* from the Canadian Atlantic Ocean, Can. J. Zool. 74 (1996) 1590–1593. https://doi.org/10.1139/z96-173.

[111] E.G. Kuznetsova, Monogenea from Chondrichthyes of the Patagonian Shelf., Ekol. Eksp. Paraziologyia. (1975) 143–153 (In russian).

[112] L.G. Poddubnaya, W. Hemmingsen, D.I. Gibson, Ultrastructural observations of the attachment organs of the monogenean Rajonchocotyle emarginata (Olsson, 1876) (Polyopisthocotylea: Hexabothriidae), a gill parasite of rays, Parasitol. Res. 115 (2016) 2285–2297. https://doi.org/10.1007/s00436-016-4973-x.

[113] G.A. Bristow, B. Berland, *Dictyocotyle coeliaca* Nybelin, 1941 (Monogenea) from the west coast of Norway, Sarsia. 73 (1988) 283–286. https://doi.org/10.1080/00364827.1988.10413414.

[114] J.W. Crane, Systematic and new species of marine Monogenea from California, Wasman J. Biol. 30 (1972) 109–166.

[115] E.W. Price, North American monogenetic trematodes. ii. The families Monocotylidae, Microbothriidae, Acanthocotylidae and Udonellidae (Capsaloidea), J. Washingt. Acad. Sci. 28 (1938) 183–198.

[116] M. Álvarez, W. Aragort, J. Leiro, M. Sanmartín, Macroparasites of five species of ray (genus *Raja)* on the northwest coast of Spain, Dis. Aquat. Organ. 70 (2006) 93–100. https://doi.org/10.3354/dao070093.

[117] M.M. Irigoitia, D.M.P. Cantatore, G.E. Delpiani, I.S. Incorvaia, A.L. Lanfranchi, J.T. Timi, *Merizocotyle euzeti* sp. n. (Monogenea: Monocotylidae) from the nasal tissue of three deep sea skates (Rajidae) in the Southwestern Atlantic Ocean, Folia Parasitol. (Praha). 61 (2014) 206–212. https://doi.org/10.14411/fp.2014.031.

[118] G. Rees, J. Llewellyn, A record of the trematode and cestode parasites of fishes from the Porcupine Bank, Irish Atlantic Slope and Irish Sea, Parasitology. 33 (1941) 390–396. https://doi.org/10.1017/S0031182000024598.

[119] E.W. Price, A redescription of *Onchocotyle emarginata* Olsson, 1876,(Trematoda: Monogenea)., Proc. Helminthol. Soc. Wash. 7 (1940) 76–78.

[120] P. Olsson, Bidrag till skandinaviens helminth fauna, K. Sven. Vetensk.-Akad. Handl. 14 (1876) 1–35.

[121] P. Sonsino, Sull’ *Octocotyle* (Vallisia) striata Par. e Per. Replica ai Parona e Perugia., Zool. Anz. 14 (1891) 87–8.

[122] P. Sonsino, Notizie di trematodi e nematodi collezione del Museo di Pisa, Atti SOCT. Osc. Xci. Nat. Pisa,p Roc. Verb. 7 (1890) 173–8.

[123] A.C. Henderson, J.J. Dunne, An introduction to the parasites of the thornback ray *Raja clavata* L. from the west coast of Ireland, Irish Natl. J. 26 (1999) 172–174.

